# *De novo* Genome Sequencing and Annotation of a Thermophilic Cyanobacterium using the Oxford Nanopore Technologies Sequencing Platform

**DOI:** 10.1101/2024.03.13.584795

**Authors:** Dirk V. Woortman, Martha Kandawa-Schulz, Norbert Mehlmer, Thomas B. Brück

## Abstract

In this project the MinION DNA sequencing platform was tested and introduced. It was proven that Next-Generation Sequencing (NGS) is not restricted to core facilities, but can be implemented in a standard laboratory setting. This makes the MinION suitable for decentralized strain screening, development and quality monitoring applications in the biotechnology industry. Over the course of the MinION early Access Program, the device was used to sequence a novel extremophile cyanobacteria candidate isolated from a hot spring at Klein Barmen, Namibia. The *de novo* assembly of the microorganism genome resulted in one bacterial chromosome without gaps. Further analysis showed, that significant variation/heterogeneity between related genomes is present. This underlines the novel character of the sequenced genome.

## Introduction

Recent advances in next-generation sequencing technology, have enabled decentralized high-throughput DNA sequencing. Oxford Nanopore Technologies™ is the first to have made a long-awaited nanopore DNA sequencer available to a broader scientific audience. The company selected a number of researchers (∼1000) for an initial beta test and developmental phase. Which was the initial foundation for this work. Being a member of the MinION™ device early access community (MAP) was a great honour.

This chapter will focus on the fundamental theoretical background in order to make the presented work comprehensible. The first section starts with DNA sequencing history, thereafter moving on to the Oxford Nanopore™ MinION™ the DNA Sequencing device. The minute sequencing platform and its performance will be presented and discussed throughout the study. At the end of this chapter the cyanobacteria phylum will be introduced.

Everything starts with DNA - essentially every living being has its fundamental properties from deoxyribonucleic acids. Nucleic acids were first isolated and identified and given a name by Friedrich Miescher in 1860.^3,4^ The structure was determined about 100 years later by Watson and Crick.^5^ In living nature two different classes of nucleic acids are found: ribonucleic acids and deoxyribonucleic acids.^6^ Deoxyribonucleic acids are built up by (deoxy) nucleotides containing four distinct bases. These are known as **A**denine, **G**uanine, **T**hymine and **C**ytosine - the building block of genomes. The research field decoding this polymer is called “Genomics”. DNA sequencing is the process of determining the order of nucleotides within a DNA molecule most accurately possible.

**Figure 2.**
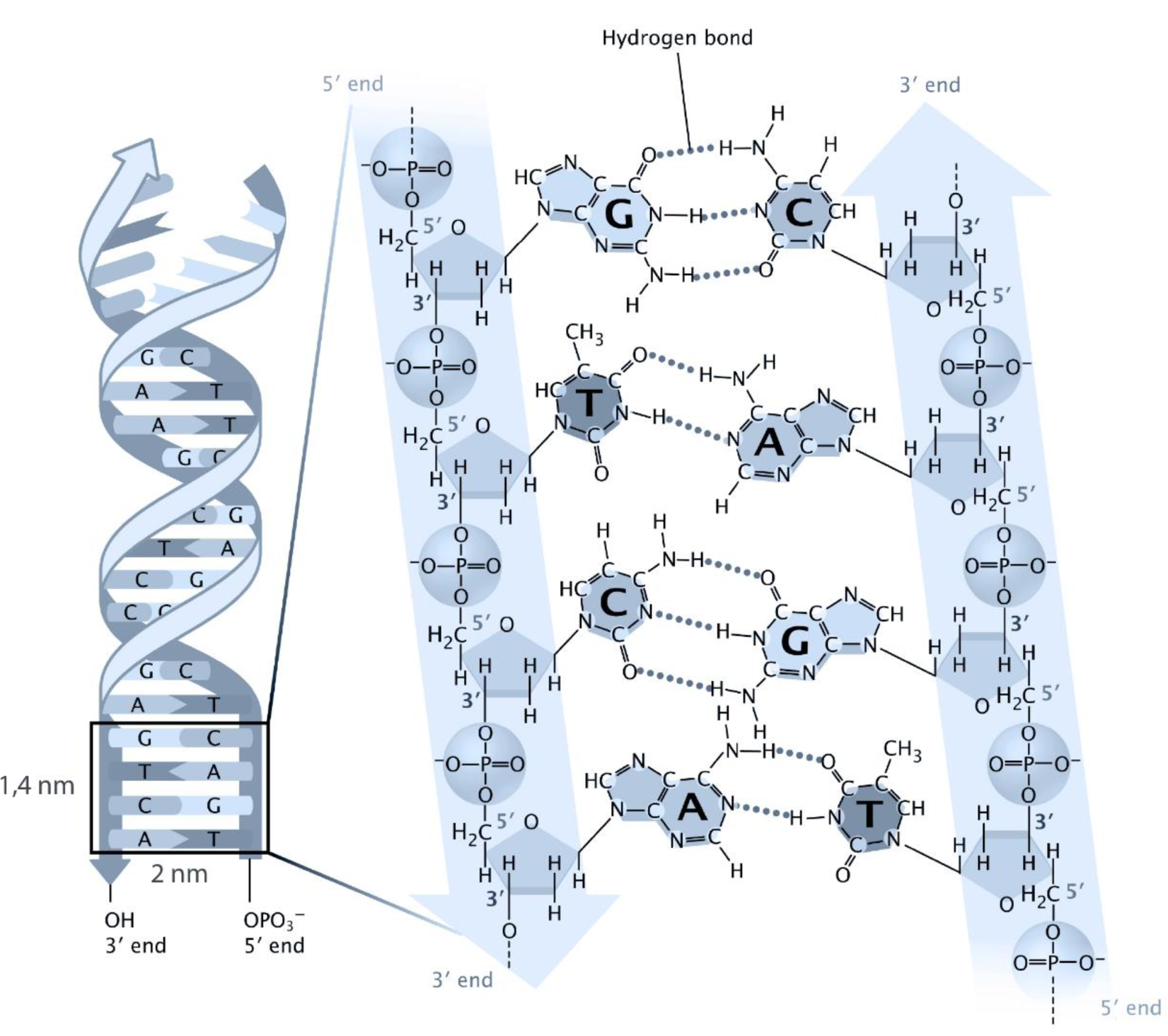
Molecular structure of DNA. Friendly provided by Jane Whitney.

During this project, only genomic DNA sequencing was studied – but it is important to note that RNA sequencing and so-called “transcriptomics” is a similar field of research with an ever-increasing application spectrum.

In figure 3 the table illustrates the differences of the building blocks of RNA and DNA. The composition of RNA and especially its secondary structure differs to DNA, and direct sequencing is very limited possible to date. Nevertheless, there are great hopes to achieve this goal in the near future by nanopore sequencing.^7^

**Figure 3.**
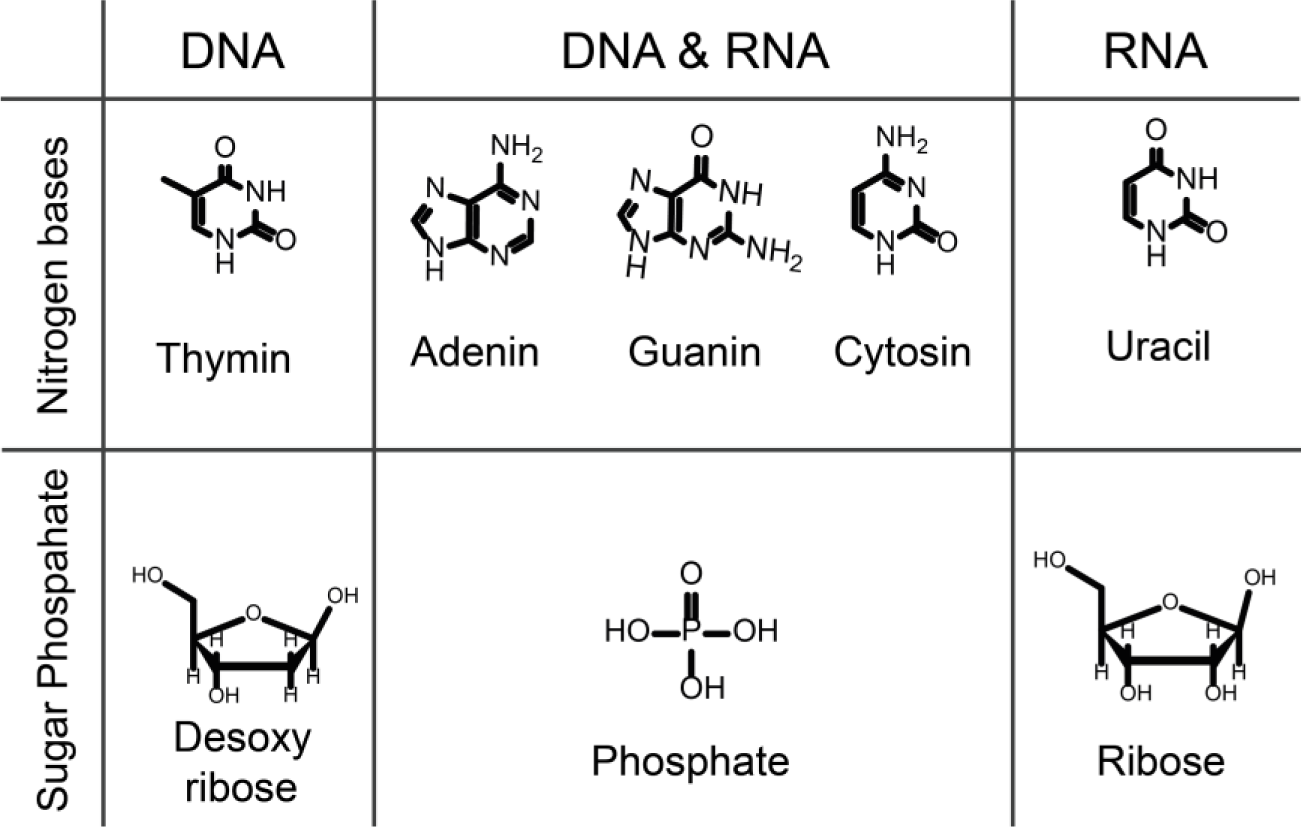
Building blocks of nucleic acids.

In the past 20 years, DNA sequencing technology evolved most rapidly - a “16-month doubling time for the sequence database is substantially faster than the 24-month doubling in computer power predicted by Moore’s law”.^8^ The best way of visualising this is by presenting the famous NIH cost diagram^9^ of DNA sequencing.

**Figure 4.**
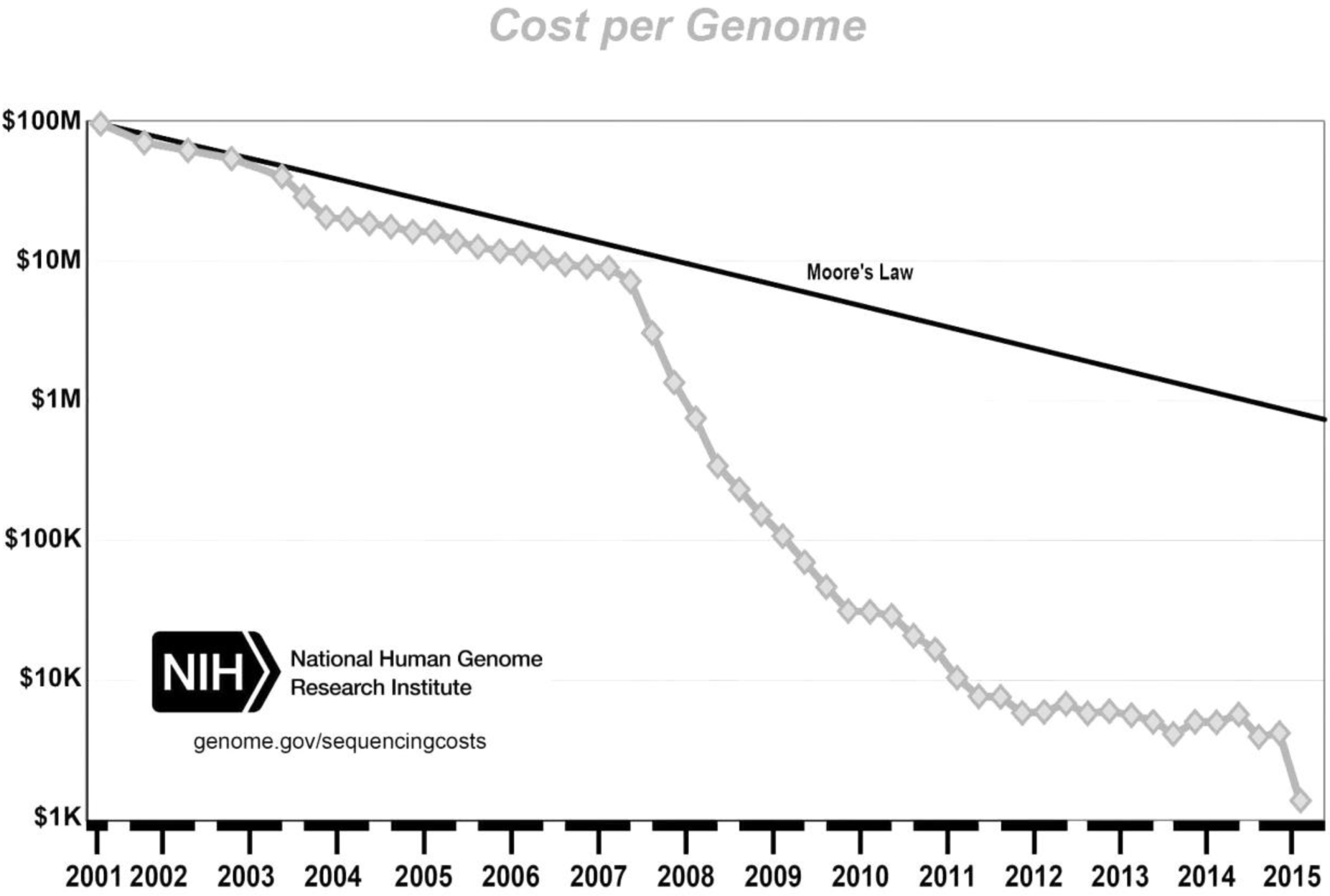
NIH diagram of DNA sequencing costs (adapted to grayscale by Adobe AI)^9^. Appearance of second-generation sequencing in ∼2007 (454)

In the meantime, DNA sequencing technologies have been categorized in the following groups by Schadt et al.:

“Sanger sequencing—the first-generation, amplification-based massively parallel sequencing—the second-generation, and single-molecule sequencing—the thirdgeneration”^7^. The popular term Next-Generation Sequencing (NGS) was introduced since the second generation sequencers were available. In the following way NGS is defined on the Nature webpage: “Next-generation sequencing refers to non-Sangerbased high-throughput DNA sequencing technologies. Millions or billions of DNA strands can be sequenced in parallel, yielding substantially more throughput and minimizing the need for the fragment-cloning methods that are often used in Sanger sequencing of genomes.”^10^

To date one can choose from a multitude of DNA sequencing platforms. In the following figure 5, an overview of the most important “NGS” companies and their milestones is shown. The most recent comparisons of sequencing technologies can be extracted from the “field guide to NGS” which is updated regularly. ^11^

**Figure 5.**
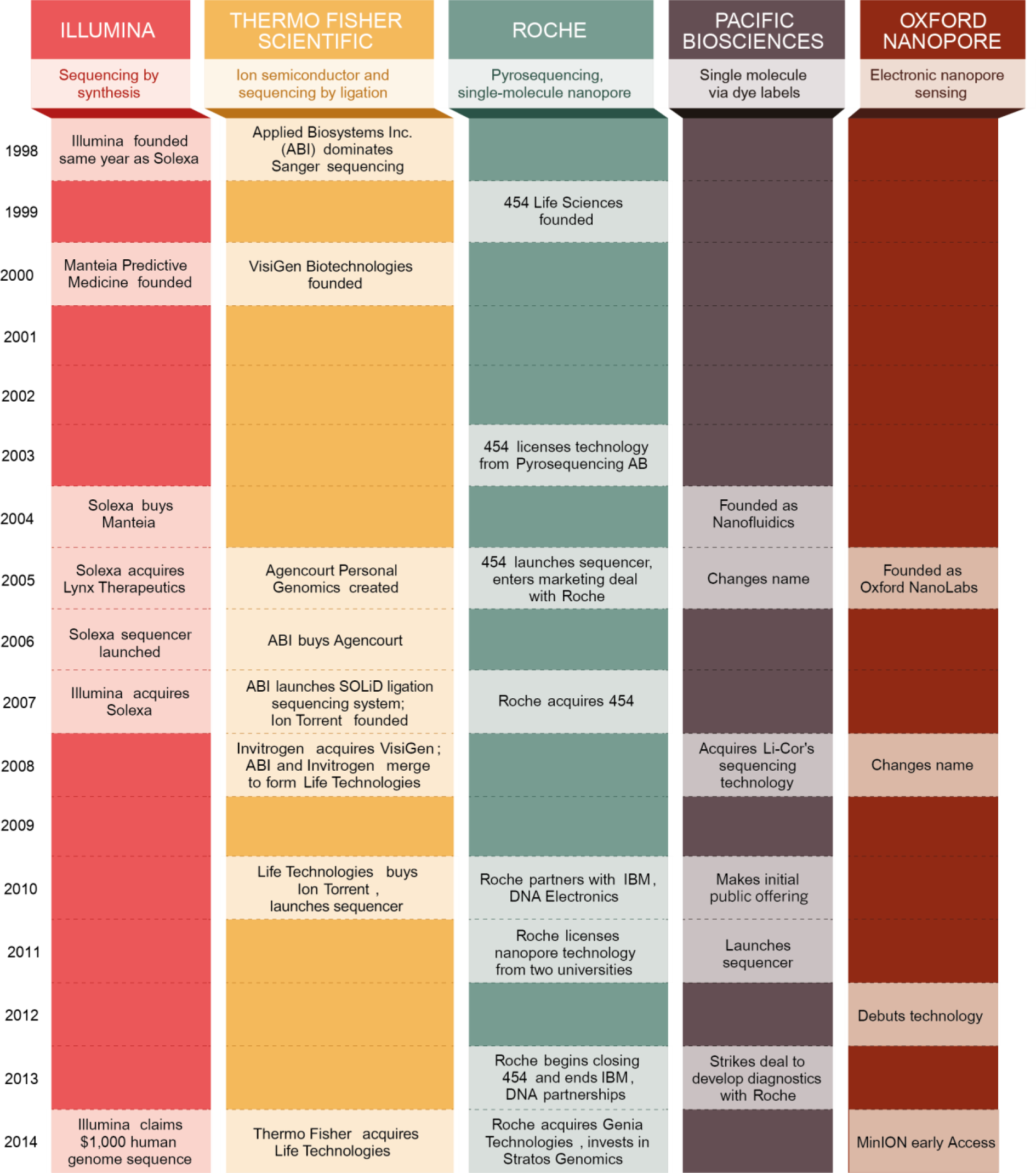
Major next-generation DNA sequencing companies and their milestones.^12^ ONT announced the MAP in April 2014.

In general, modern genomics technologies rely on three common steps: “template preparation, sequencing and data analysis.”^13^ For all current sequencing technologies DNA has to be isolated and fragmented first. Depending on the sequencing technology used, the DNA is enzymatically/chemically or physically sheared into appropriate sizes.^13^ The sizes vary currently from 20 to 100,000 bases.^14^

As microfluidics handling for routine applications gets problematic with DNA fragment sizes near 100 kb ^9^ an optimum can be expected somewhere in this range, in the near future.

In the following step the so-called “library” is prepared. That means fragmented gDNA is transformed into a form, which can be analysed by the sequencing technology. The term “library” refers to modified gDNA fragments which can be loaded on the sequencing device. Thereafter the library is “loaded” on the system and will be sequenced. The data can be analysed and interpreted after termination of the sequencing run, with runtimes varying from two hours to 14 days.^11^ The result of the sequencing run is a large collection of reads saved in FASTA/FASTQ or a specific file format. Finally, the data can be obtained and used to reconstruct a model of the organism’s genome.

The bottom up genome analysis^15^ includes a reads filtering step followed by a genome (re-)assembly process. The results can then be analysed, compared and fed into databases.

**Figure 6.**
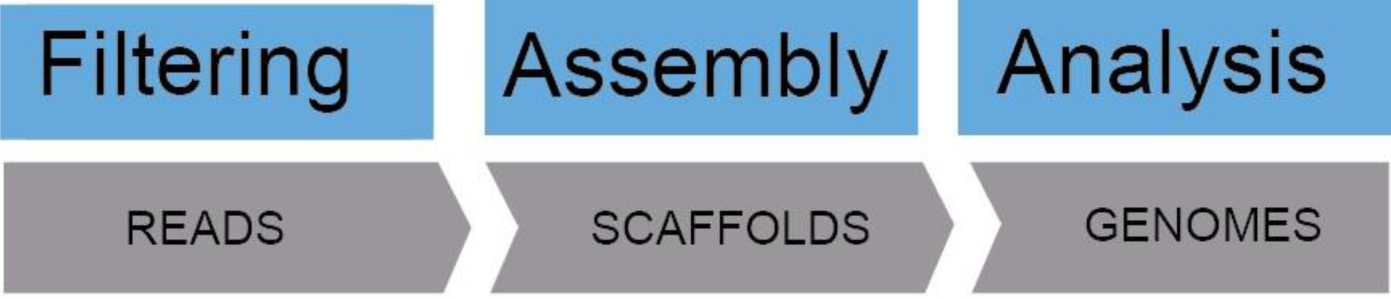
*De novo* genome analysis workflow.

The workflow includes an initial read QC filtering step which importance is often overlooked^16^. During the filtering, the raw data is reviewed and judged for the first time. Based on their quality the reads are then trimmed and subsets are generated. At this point, depending on data yield and quality, one can decide to carry on accumulating sequencing data or move on to the assembling process.^1^ When the data yield is sufficient the next step is *de novo* genome assembly. Assembling the reads to a full *de novo* genome is a complex bioinformatics task.^17^ Keith Bradnam (UC Davis) gave the following definition while highlighting the word attempt: “A genome assembly is an attempt to accurately represent an entire genome sequence from a large set of very short DNA sequences.”^18^

Since the Human genome sequencing project - not much has changed regarding a whole genome assembly process. The task is carried out by a genome assembler, software which assembles the reads to genomes, by using an algorithm which finds overlap regions of all reads. There are more than 125 assemblers one can choose from and each assembler has multiple command line options.^18^ In principle the assembler firstly constructs consensus sequences from the reads – so-called contigs. These are further aligned to form scaffolds (at best only one), which represent the genome.

The Celera whole-genome shotgun (WGS) DNA sequence assembler is one of such software packages. A lot of literature can be found about this assembler as it was developed for the Human genome project.^19^ Moreover, the Celera assembler can handle long MinION™ (which will be introduced later on) reads.^2^

As the MinION™ data is error prone (15%)^20^ some adaptions to the established assembly processes had to be made.^2^ These adaptions to the assembly process were a bioinformatics breakthrough enabling MinION™ and PacBio reads to be successfully assembled.^21^ These adaptions resulted in the hierarchical genome assembly process (HGAP).^22^ This process includes a reads correction step, thereafter the data is assembled by a standard assembler, a final polishing step completes the HGAP.

The read correction step uses the “longest reads as seeds, recruiting all other reads to construct highly accurate preassembled reads (using a direct acyclic graph-based consensus procedure)”^22^. These (preassembled) corrected reads are then assembled by Celera to form the draft genome. Thereafter, the draft genome is polished with the initial raw data once again.^22^ In figure 7 this process (except the polishing) is visualized by the original publicized diagram.

**Figure 7.**
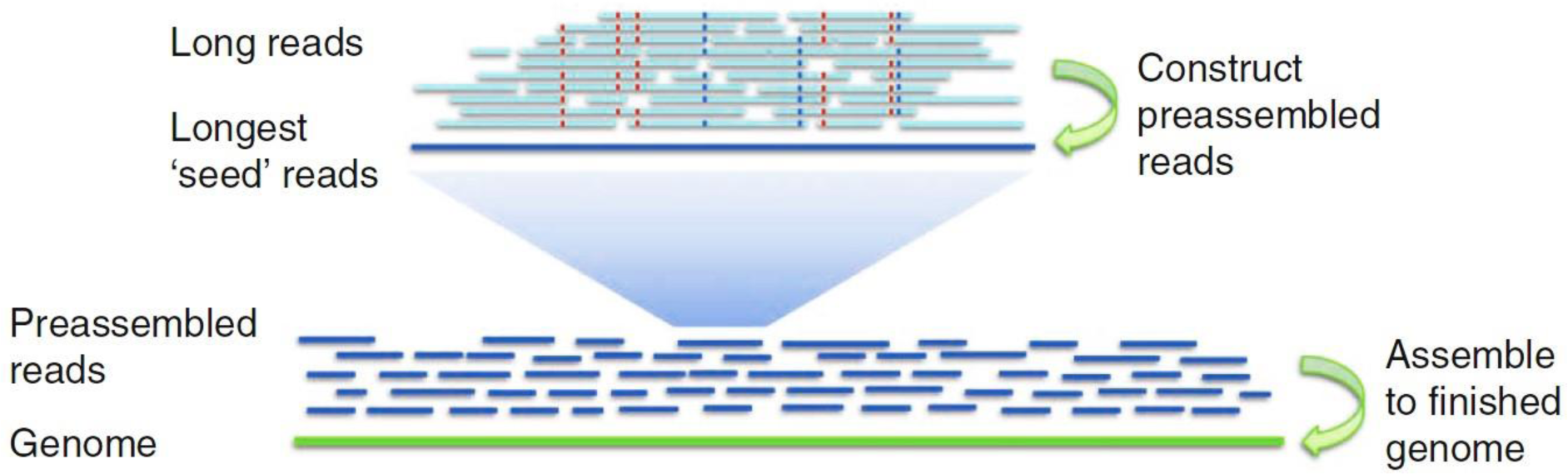
*De novo* assembly process using the HGAP (hierarchical genome assembly process)^22^.

In accordance to the HGAP procedure, a *de novo* assembly process was suggested for MinION™ data by Loman et al. The proven concept from Loman et al. (*E.coli*) was used to assembly the genome sequenced during this work.

To complete this introduction a list of tools and their algorithms used for correction, assembly and polishing are presented in table 1.

**Table 1.** Tools used for the assembly process.

| Process | Tool | Algorithm |
| --- | --- | --- |
| Correction | Nanocorrect<sup>2</sup> | Partial order alignment<sup>23</sup> |
| Assembly | Celera | Greedy-Path-MergingAlgorithm<sup>24</sup> |
| Polishing | Nanopolish | HMM / POA<sup>23</sup> |

Once the filtered and pre-processed reads are assembled and polished the final genome can be analysed by further bioinformatics software. The assignment of functions to the *de novo* genome is done by annotation software using a number of databases to predict possible proteins, regulative regions and pathways.

### DNA Sequencing with Oxford Nanopore Technologies™ MinION™ device

Oxford Nanopore Technologies™, initially started from academia,^25^ has recently made the first nanopore sequencer available. The MinION™ device, being smaller than a modern mobile telephone, introduced a new era of portable DNA sequencing^26^ regarded by Nicholas Loman as the “democratisation of sequencing”. Furthermore, it has been shown by a NASA (National Aeronautics and Space Administration) team that DNA sequencing can be done in low gravity environment.^27^ The “in space” sequencing project had been announced during Oxford Nanopore Technologies™ first “London Calling” Conference.

### The principle of nanopore sequencing and its origin

Nanopores are tiny holes through any material with diameters in the nanometre scale. The core of the Oxford Nanopore™ sequencing technology is a channel forming protein.^1^ Channel proteins have been known and studied for a long time in the field of electrophysiology. Erwin Neher (* 20. March 1944 Landsberg am Lech, Bavaria) as well as Bert Sakmann (* 12. June 1942 Stuttgart) were awarded 1991 a Nobel prize for the Patch Clamp Amplifier method (PCA)^28^. This resulted in a long journey of research in measuring currents flowing to tiny holes – known as nanopores. The rapid appearance of the term nanopore in scientific literature is shown in figure 8.

**Figure 8.**
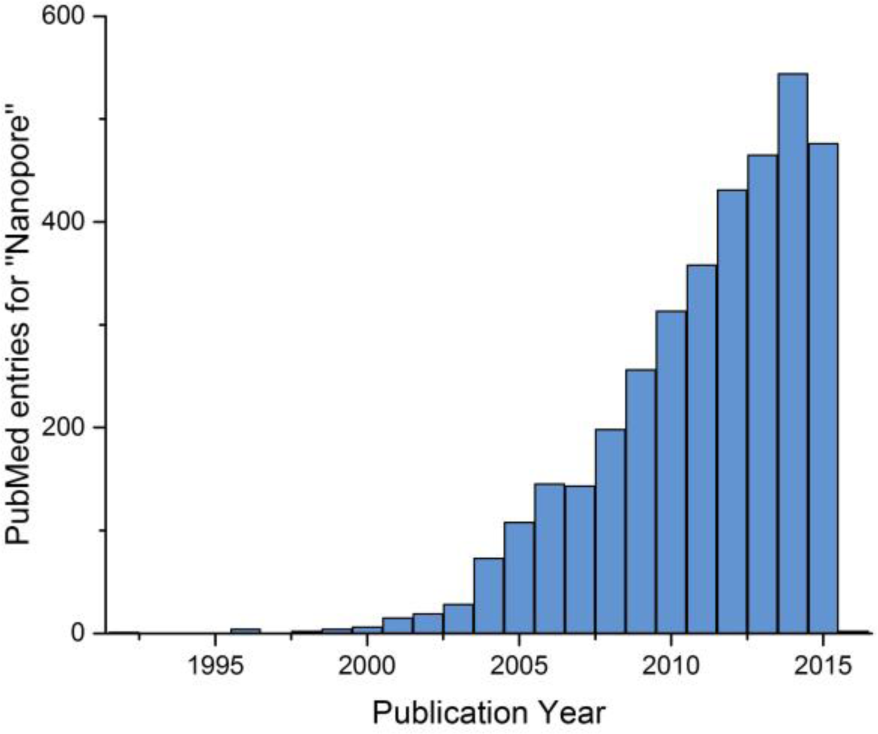
“Nanopore” appearance in the primary literature. Graph plotted with Origin Pro, data extracted from PubMed 25.11.2015.

### The Principle

Nanopore sequencing is a relative old sequencing concept – about 25 years ago it was mentioned the first time.^25^ The technology has relied on many different advancements in the field of microbiology, genetics, electronics and microfluidics as well as informatics.

The Oxford Nanopore™ sequencing technology relies on the interaction of channel proteins and an intact DNA strand. This process can be measured directly and in real time within the MinION™ flow cell.^1^ To measure the “interaction” a “constant” electronic potential of about −150 mV is applied over a nanopore embedded membrane, resulting in a measurable current (∼ 10^2^ pA (1 pA = 6.25 × 10^6^ charges per second)^25^) flowing through the pores. While this is happening, a DNA strand can be propagating through the minute channel creating distinct fluctuations and disruptions in the current.^1^ This concept can be seen below in figure 9.

**Figure 9.**
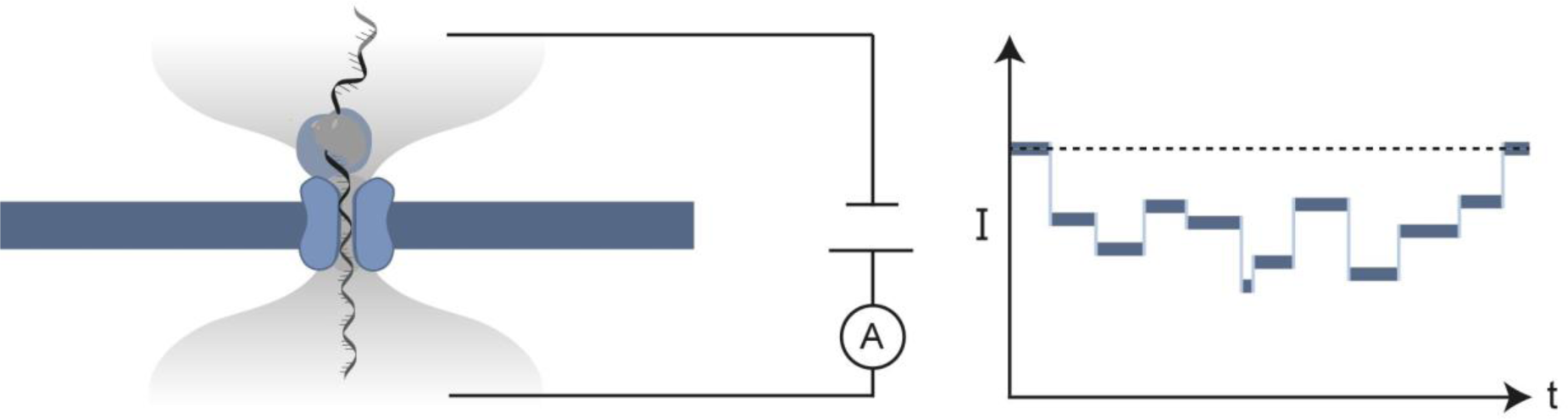
The principle of nanopore DNA sequencing.

### The Origin

The origin of nanopore sequencing dates back to approximately 1996 when Kasianowicz et al. first measured the translocation of DNA through a nanopore^29^. At that time, he could measure the time needed for a DNA fragment to translocate. He postulated in 1996 that “with further improvements, the method could in principle provide direct, high-speed detection of the sequence of bases in single molecules of DNA or RNA”.

Single base resolution is a fundamental requirement for DNA sequencing and therefore was a sought-after achievement.^7^ Major improvements towards single base identification were brought to this technology by Mark Akeson et al. They found a setup where a “Nanopore behaves as a detector that can rapidly discriminate between pyrimidine and purine segments”.^31^ The protein pore (seen in figure 10) used by Akeson et al. was derived from α-haemolysin – a *Staphylococcus aureus* bacterial toxin known for its lysis of erythrocytes by poration of the cell membrane. The pore is a heptamere which has some complex membrane associated assembly pathways from its monomers.^32^

**Figure 10.**
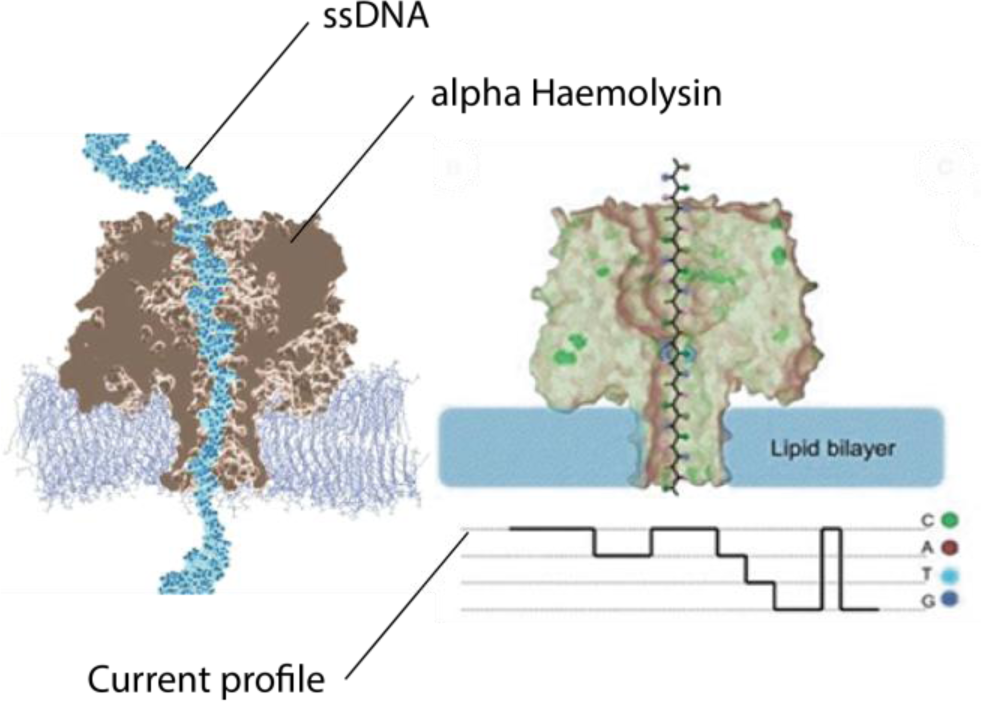
MD simulation of the 232 kDa Alpha-Haemolysin with ssDNA from Klaus Schulten et al.^30^ and to the right an illustration of the pore and the resulting current profile from Wang ^7^

The narrow 1,4 nm diameter congestion lets only single stranded DNA (ssDNA) progress through the pore.^33^ In figure 11 the α-haemolysin congestion is shown on the left. Furthermore, real residual current recordings of single base translocations can be seen in the centre and the corresponding residual pore current distributions on the right.

**Figure 11.**
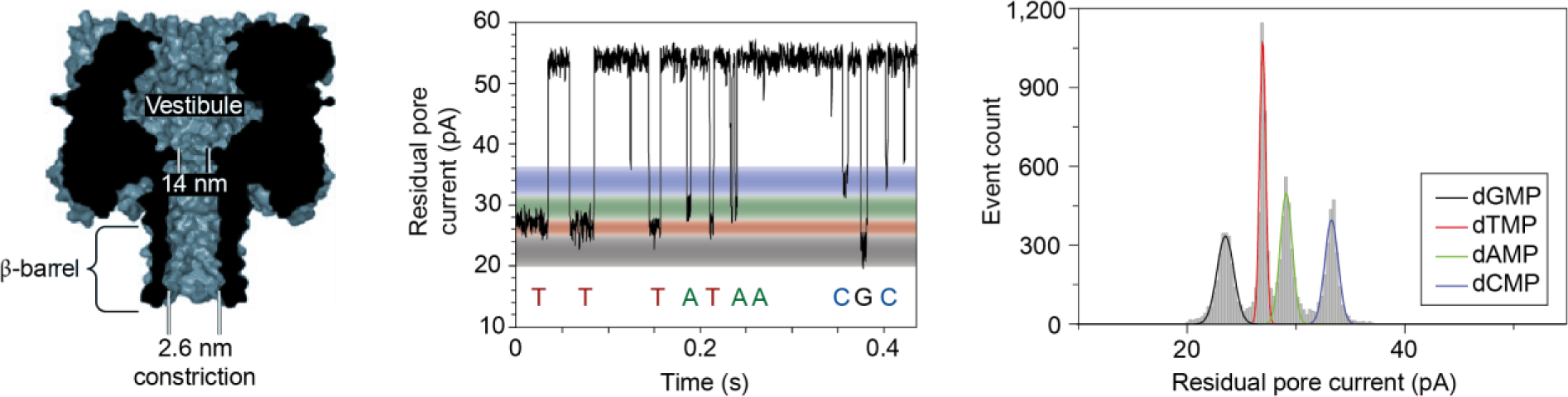
α-haemolysin and its ability to distinguish between nucleotides. From left to right: The αhaemolysin dimension. Characteristic current disruptions for each single base. Distribution of the specific residual current for the “bases”. ^34^

An important breakthrough was slowing down the translocation process. In this field a lot had been achieved by Bayley et al. who studied the recognition sites.^35^ By targeted addition of cyclodextrins further improvements and different specificity were demonstrated.^36^ Hagan Bayley extensively studied α-haemolysin and is an intellectual property (IP) holder of ONT.^25^ The exact identity of the Oxford Nanopore Technologies™ protein-pore, used within the MinION™ technology is not clear^37^. But it is clearly an optimisation problem with a wide sample space.

### The Oxford Nanopore™ sequencing technology

Oxford Nanopore Technologies™ DNA sequencing method, relies on the progression of single stranded DNA through a protein pore. Using a protein as nanopore has distinct production, sensing and engineering advantages.^34^

In the next passage the “chemistry” used by Oxford Nanopore Technologies™ will be introduced. This part is generally called “Wetlab”. Thereafter, introducing the hardware and software of the MinION™ device – called the “Drylab”.

### The Wetlab

The so-called “chemistry” is fundamental to the controlled translocation of DNA. The technological breakthrough was to slow down the process of DNA translocation by adopting a motor-protein^38^ to “ratchet” the **ss**DNA with a “constant” speed through the pore.^39^ The motor-protein is an “ATP-fuelled ratcheting enzyme”, similar to a helicase which unwinds **ds**DNA in a controlled manner.^40^

Starting from one end the template **ss**DNA is translocated through the pore by ∼30 bases per second. The complement strand stays (first of all) in the library compartment.^1^ A technology to sequence both strands of the **ds**DNA molecule in one continuous movement is patented by ONT^41^. The idea of ligating a hairpin adaptor to one of the **ds**DNA ends was secured. This essentially makes one continuous **ss**DNA molecule from a Watson-Crick paired (double stranded) DNA molecule. The connection of both strands (by the hairpin) makes it possible for the motor-protein to “carry on” translocating the complement strand as soon as the hairpin is reached.^1^ In figure 12 the concept is visualized.

**Figure 12.**
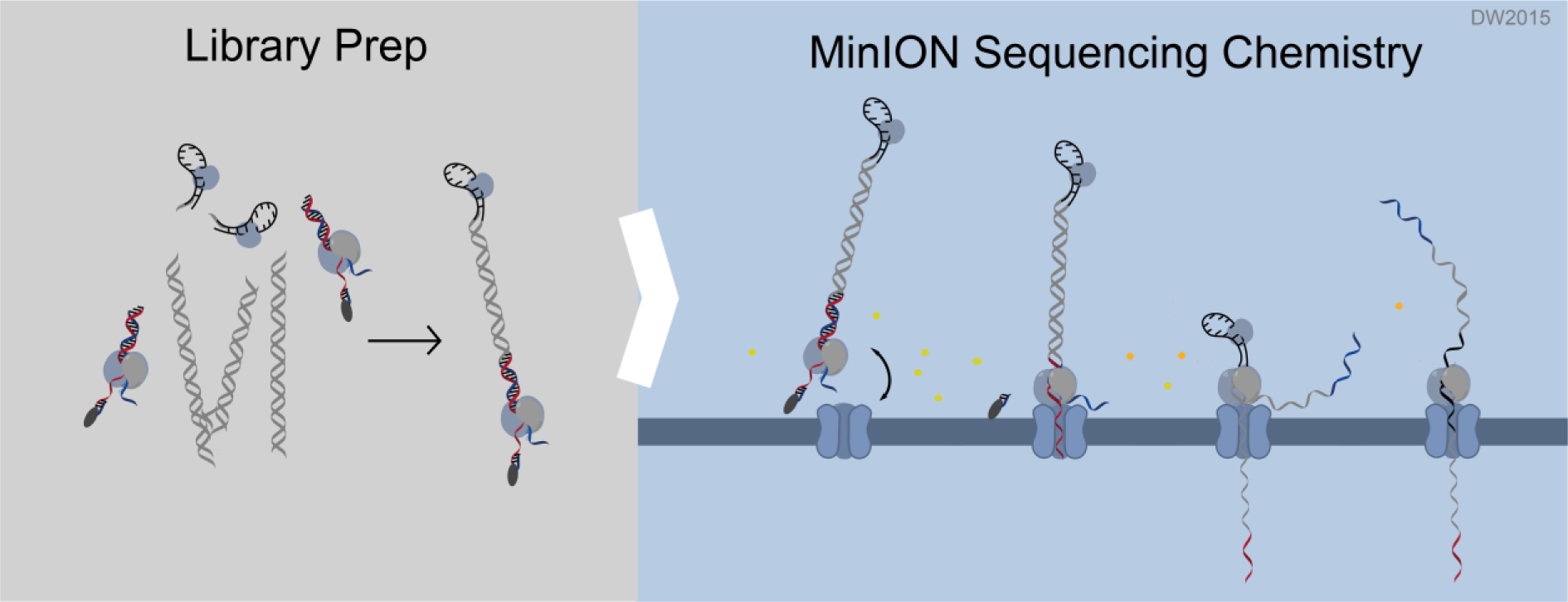
The MinION™ technology. Left: Ligation of adaptors (AMX HPA(black)) with the gDNA fragments. Right: Illustration of the sequencing process within the flowcell.

During library preparation two different adaptors are ligated by random chance to the DNA fragment. The first adaptor (AMX) is a dsDNA molecule with the motor protein bound. The second adaptor is a hairpin structured ssDNA molecule with a further protein bound. This protein creates a distinct pore current signal^1^, and secondly protects the bound strand from abrupt damaging forces.To the adaptors a tethering complex is bound which interacts with the nanopore embedded membrane.^20^

Once both connected strands translocated through the pore, a final 2D basecalled consensus sequence can be calculated from here on. This process is the beginning of the Drylab.

### The Drylab

To measure the process of DNA passage, ONT has evolved from the PCA (Patch Clamp Amplifiers) to very specific and sensitive ASIC (application specific integrated circuit) sensors. The drylab starts as soon as the ASIC records the current signals. The ASIC makes current readouts in more than 3000Hz - the basis for “stochastic sensing”.^25,1^ In figure 13 the raw signals (pico Ampere) (b) recorded by the ASIC while the library DNA is translocating (a) is shown. Distinct disruptions (c) can be observed for strand incorporation (i) and when reaching the hairpin (v).

**Figure 13.**
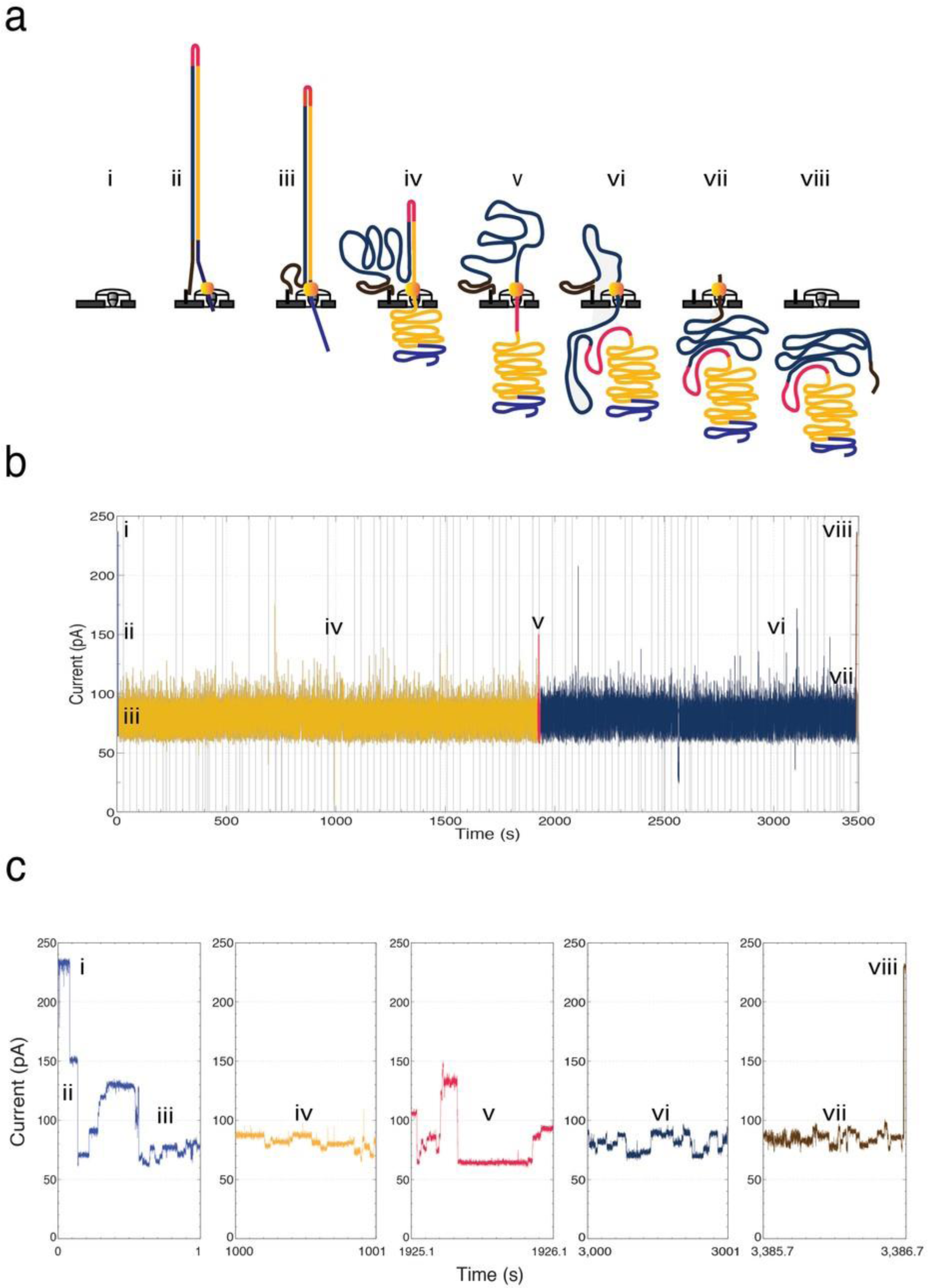
Raw signals corresponding to the different phases(i-viii) of stand progression.^42^ Illustrations by Akeson et al.

The first local computed step is the reduction of the raw signal to an abstract, more easily to interpret sequence of events. Each signal, probably corresponding to a base, is described by so-called “events”. One current disruption “event” is described by a signal mean, duration/length, standard deviation and start time, as shown in figure 14.

**Figure 14.**
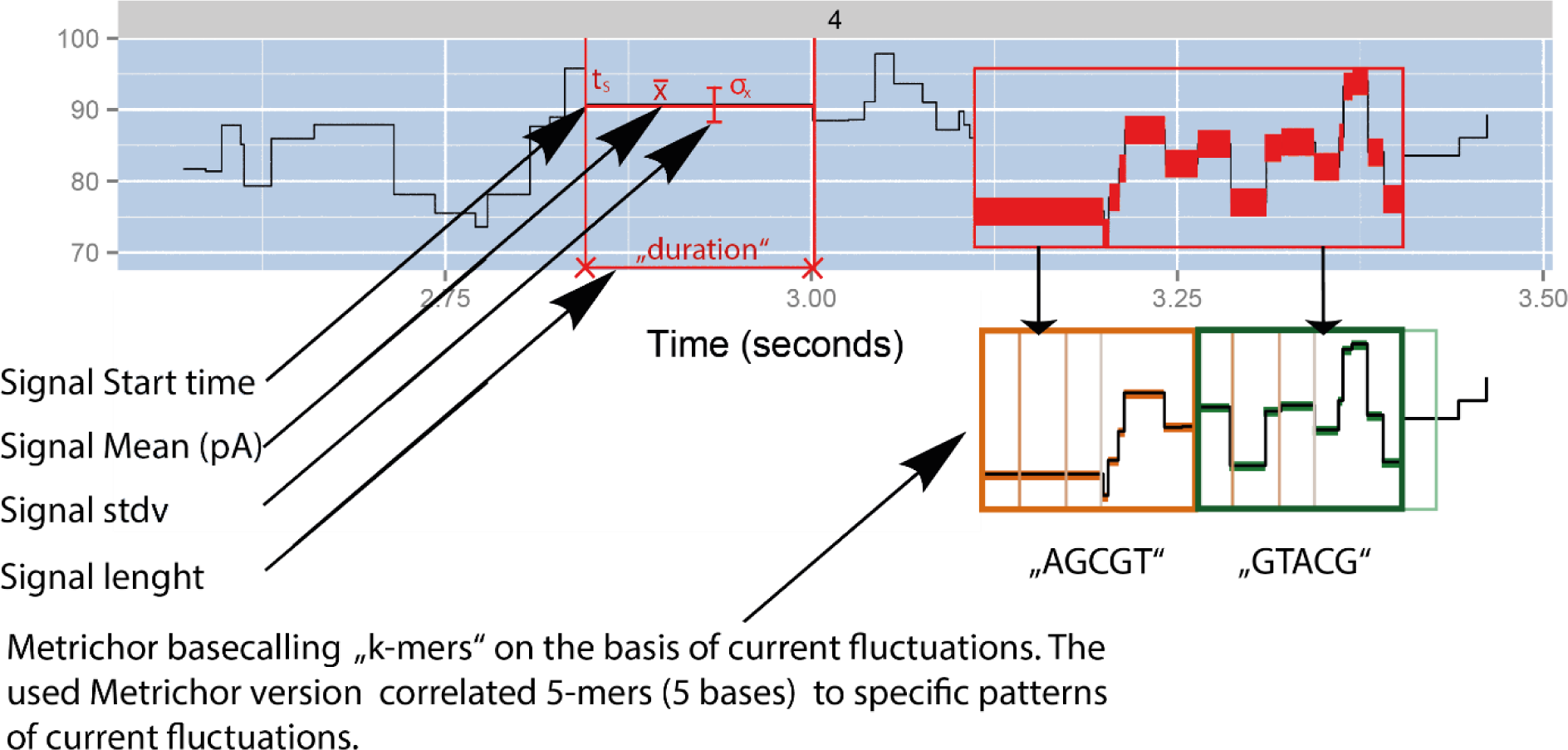
Modified squiggle plot extracted from a raw fast5 file with poretools. The parameters of the signal are included in the HDF file format additional to all FASTA, FASTq and metrics data. The read “squiggle plot” can be found in the appendix figure 43.

These “squiggle data” signals are send to an online basecalling service called “Metrichor”^43^. Metrichor uses a complex HMM/Viterbi algorithm for basecalling.^20^ It has been observed that ∼5 bases interact with α-haemolysin, thus contributing to the resulting signal.^44,35^ Therefore, Metrichor is using a 5mer model with 4^5^ (1024) distinct possible signal paths.^20^ The algorithm is “finding an optimal path of successive 5mers”^20^ in the event data model. These successive paths are “translated” into a DNA sequence.

The template/leading as well as the complement/lagging strand is basecalled independently, and by alignment a “synthetic” consensus 2D read is generated. The output is saved including all metrics and quality values in a HDF type FAST5 file. From the FAST5 file the corresponding FASTA/Q file can be extracted. As this happens in real time live analysis workflows like WIMP (What is in my pot) can be implemented.^1^

### The MinION™ device

The MinION™ device is smaller than a Mars® chocolate bar, powered by a USB cable and ultimately portable. No other DNA sequencing device to date can compete with the portability and independence from auxiliary power supply.^20^ A single use flow cell fits into the device as seen in the centre of figure 15.

**Figure 15.**
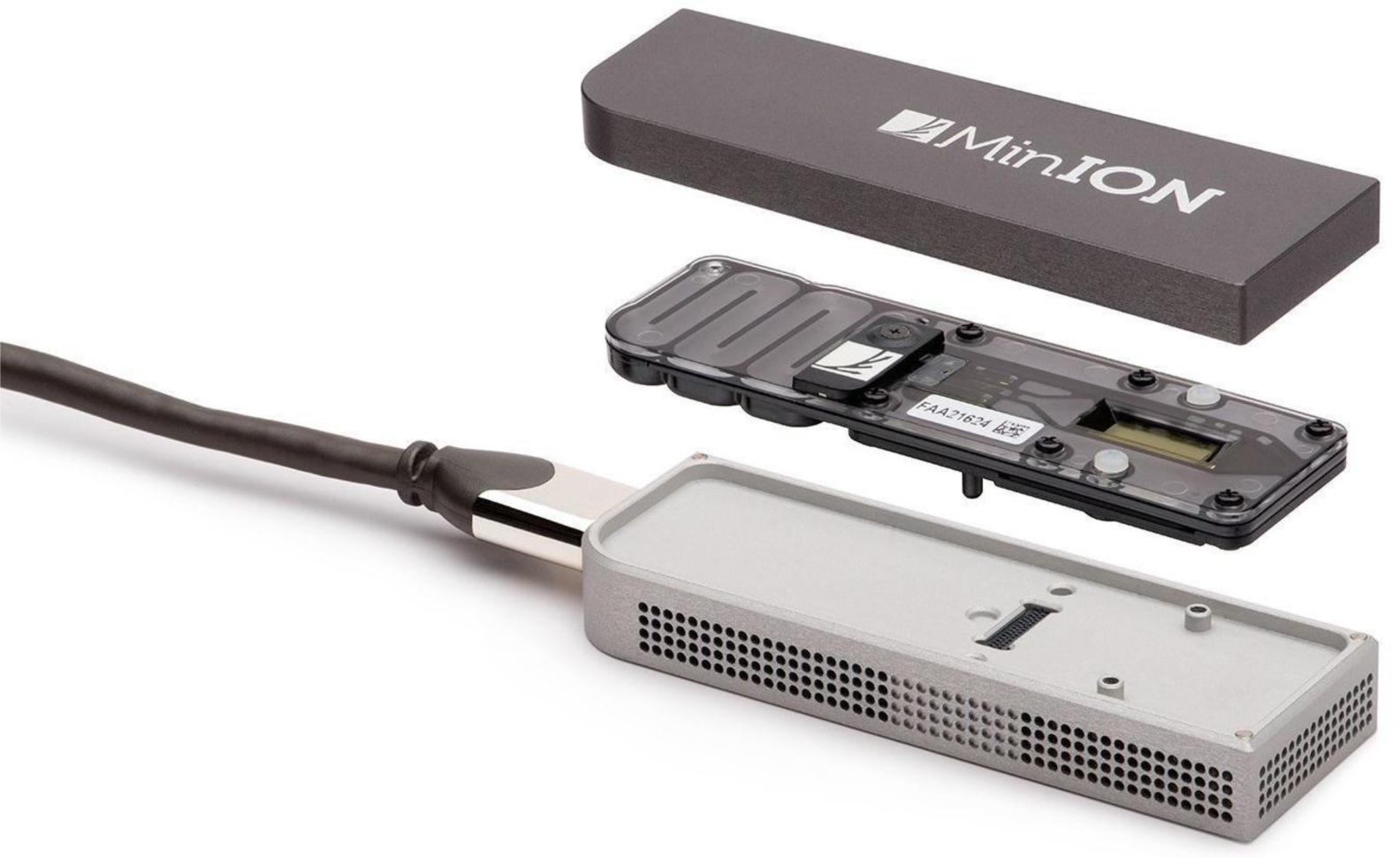
The MinION™ device in an exploded view.^1^ The flowcell can be seen in the centre. On top is a removable lid. The bottom part is showing the USB 3.0 powered device.

The flow cell has 2048 protein-pores imbedded in a membrane. These are grouped in 4 to form 512 parallel accessible sensors. The quality of the flowcell is checked before running the experiment. The process takes about 15 minutes. At the moment the flow cell is not a full disposable part, rather a recycle item. Future iteratiońs will be fully disposable with low residual value.^37^ A heatpad has to be installed for optimal temperature control during the sequencing run. The sensor array can be found in the hollow yellowish part of the flow cell in figure 15.

Derived from the provided protocol a workflow guided checklist protocol was devolved as shown in figure 16. The MinION™ is placed as a “consumer product” where laboratory staff with little NGS experience could be the customer. The idea that “untrained” researchers from adjacent fields with little NGS experience can apply the full process of whole genome sequencing was proven in this study.

**Figure 16.**
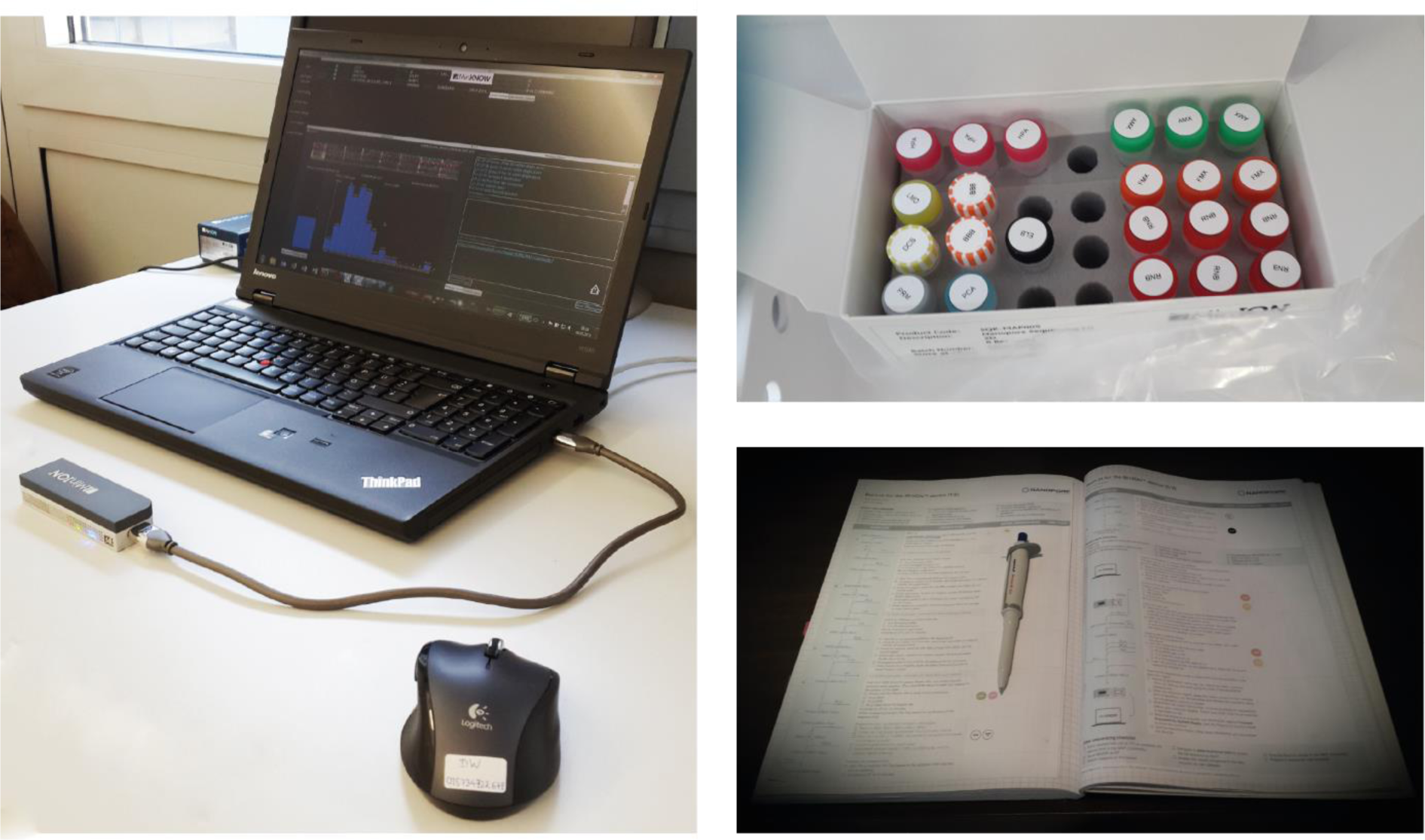
The Oxford Nanopore Technologies™ MinION™ device. Left: Workstation with handheld DNA sequencing platform. Right top: Components of the ONT Sequencing Kit for the library prep. Right bottom: Lab journal with a paper based checklist for optimal documentation and error reduced Library preparation. A tool which was developed together with ONT during the study.

### The Cyanobacteria

Cyanobacteria or blue green algae are a large diverse family of prokaryotes which are able to reduce CO2 using light, and thereby being independent from other carbons sources.^45^ They are regarded as one of the oldest living forms on earth existing more than 2500 million years.^45^ Their prokaryotic character makes them a very special group of phototropic organisms. Figure 17 puts cyanobacteria in context to the eubacteria domain. Their common name blue green algae was derived from their sometimes slight blue appearance, due to the distinct blue absorption gap shown in figure 18.

**Figure 17.**
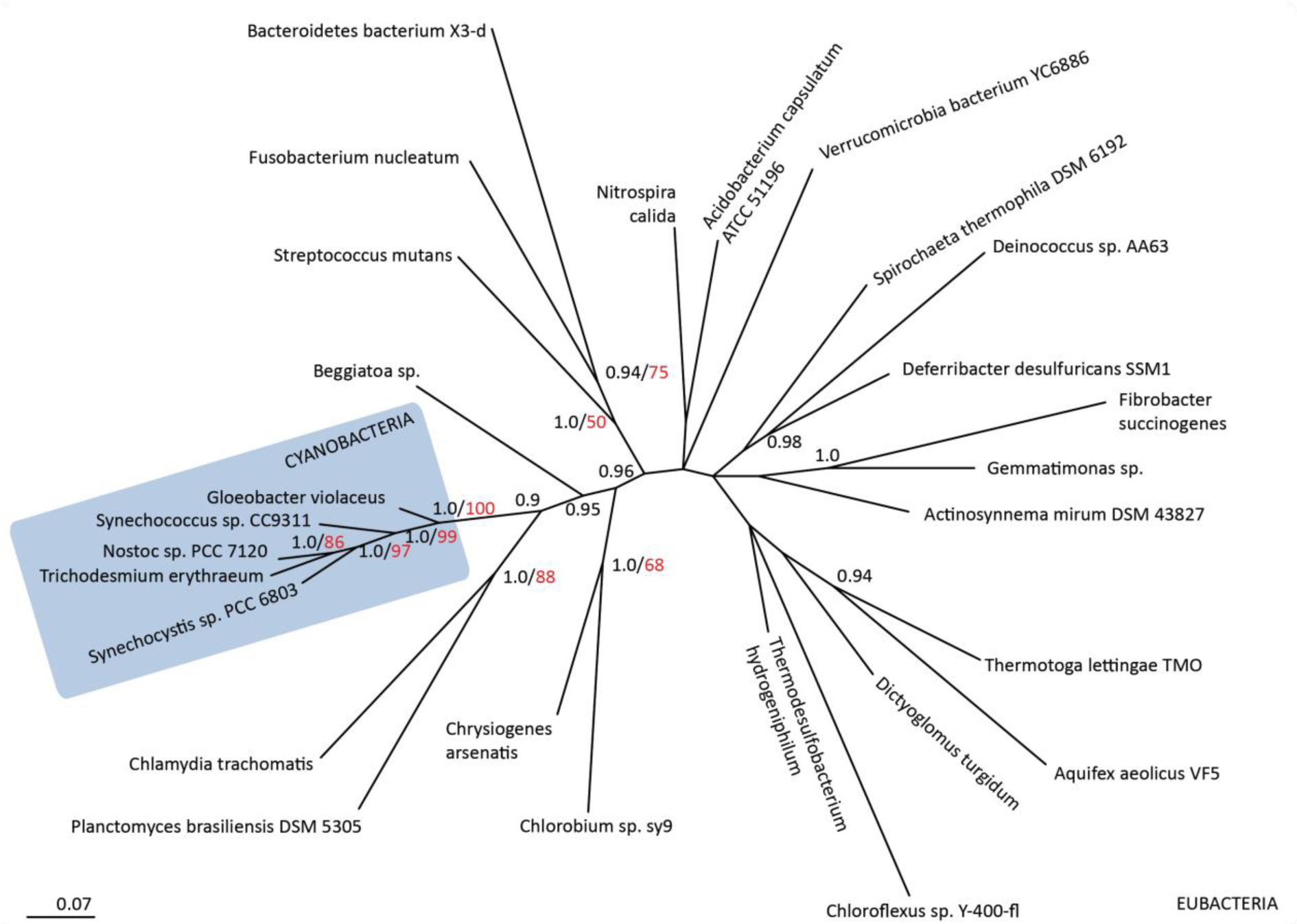
Phylogenetic tree of bacteria on 16s RNA basis. Cyanobacteria in the dark square.^46^ Modified with Adobe AI.

**Figure 18.**
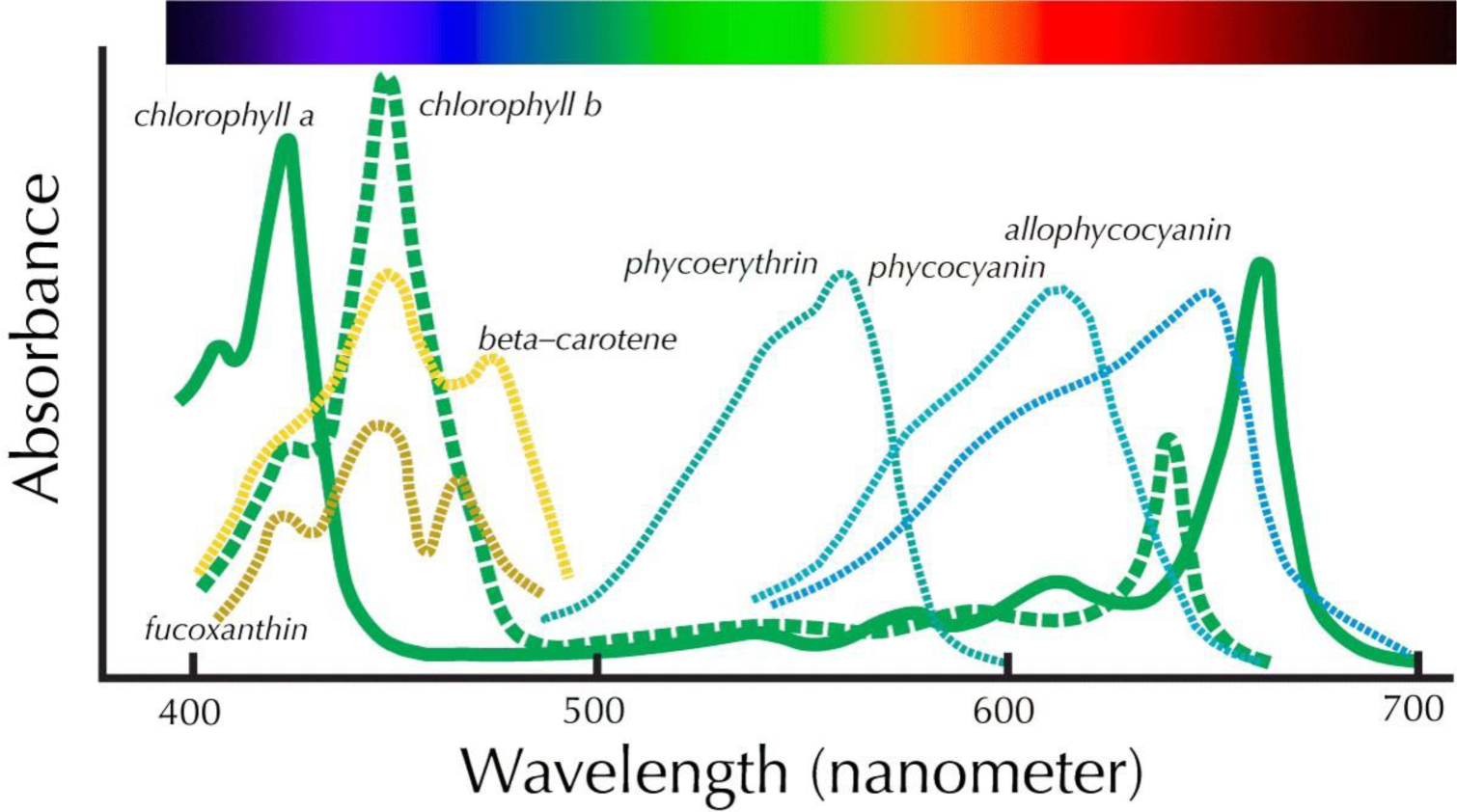
Absorptions spectra of components of cyanobacteria.^48^ Absorption gap at ∼500nm. The blue line represents pigments only associated with cyanobacterial phycobilisomes only.

There are 2 different classes of oxygenic autotroph prokaryotes, namely cyanobacteria and prochlorophyta. These are the only two groups of bacteria which carry out oxygenic photosynthesis with water as their source of electrons.^47^ These groups of organisms are distinguishable through their photosynthetic apparatus and their pigments.

Cyanobacteria produce chlorophyll a and pycobillisoms where as prochlorophyta chlorophyll a and b. From here on only cyanobacteria will be further described.

The photosynthetically active complexes are evolutionary highly conserved. (Glazer, 1983). Cyanobacteria are evolutionary decedents of the chloroplasts. In order to study the photosynthetic mechanisms, cyanobacteria have been extensively described in basic research in the past. Their transformability make them a well researchable^49^ model organism. The best studied model organism for cyanobacterial basic research are members of the *Synechocystis* and *Synechococcus* order.^50, 51^

Over the course of this study a new member of *Synechococcus* was isolated, identified, purified and extensively described by next-generation sequencing.

### Taxonomic classification

It appears to be especially difficult to systematically subdivide different groups of the cyanobacteria family.^47^ Cyanobacteria were traditionally grouped by classical botanical systems with help of uncultured environmental samples. Roughly 1500 species are believed to exist.^52^ Since many pure cultures exist by now, these have been grouped accordingly.^47^ With help of NGS it is possible to compare cyanobacteria strains within different ecosystems and shed light on still unresolved evolutionary questions.

Within the cyanobacteria family there are 5 different distinctive classes, which were subdivided by Rippka et al. in traditional botanical groups. More recent, Giovanonni et al. applied molecular phylogenetics ^47^ supporting only parts of the classic subdivisions.^45^ In figure 19 a phylogenetic overview of species characterized with help genomics is shown.

**Figure 19.**
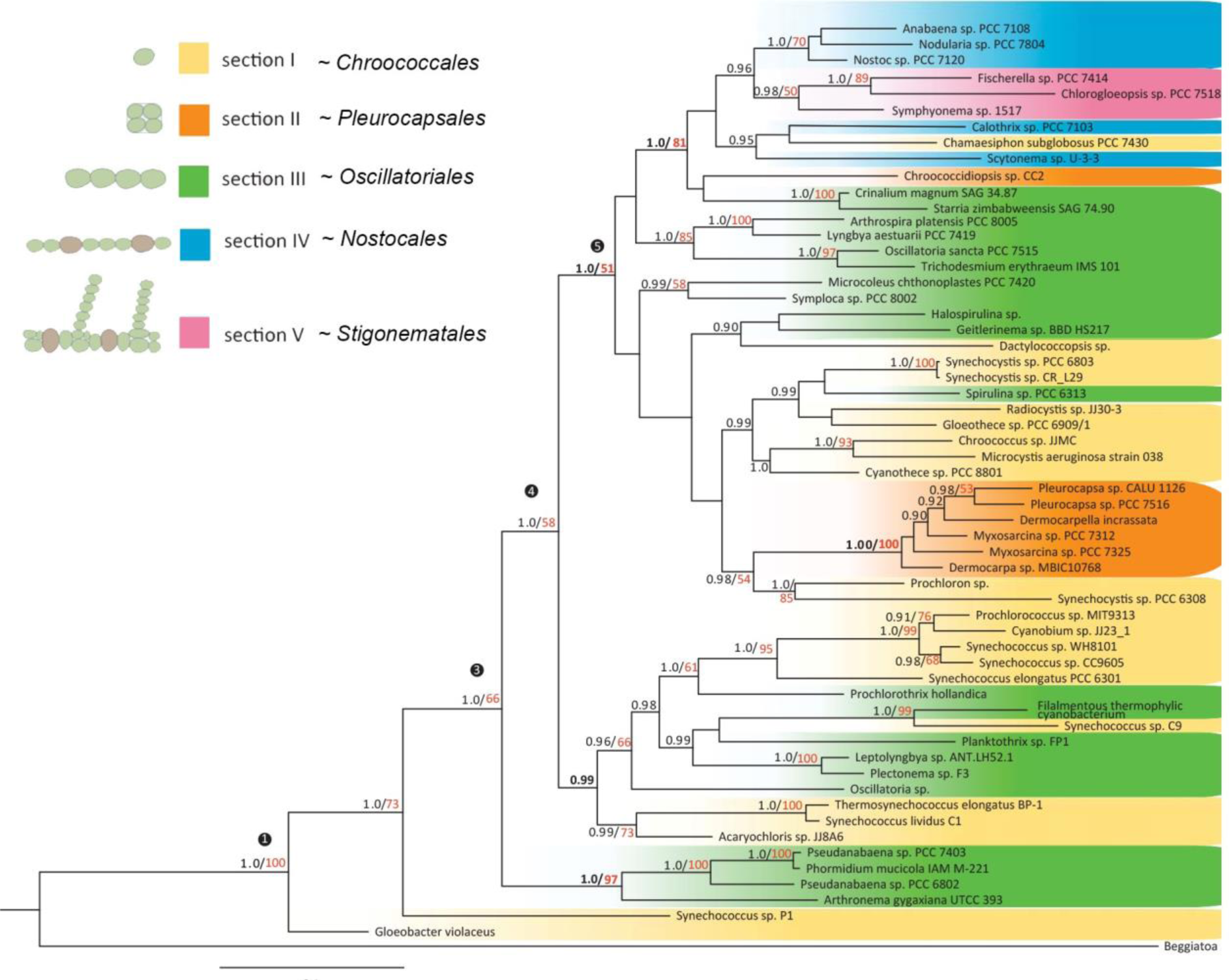
Phylogenetic tree of sequenced cyanobacteria (modified with Adobe AI).^46^.

### Thermophilic Synechococcus

The cyanobacteria sequenced during this work belongs to the thermophilic *Synechococcus* group. Thermophiles are a group of extremophiles which do grow and thrive at >45°C.^53^ Two *Synechococcus* strains which do belong to the thermophilic group have been whole genome sequenced in the past.

#### Thermosynechococcus elongatus BP-1

was one of the first cyanobacteria characterized by full genome sequencing. It was isolated in Japan from a hot spring near the city of Beppu with a growth optimum at 55°C. It has a circular genome with 2,593,857 bp, a GC content of 53,9% and no plasmids were found.^52^ An overall coverage of 5x. 2475 potential protein encoding genes could be identified in the year 2002. Some thermophilic characters were postulated including the “lack of desaturases and the presence of more heat shock proteins” than typical. This strain was the first thermophilic cyanobacteria to be transformed with exogenous DNA.^52^

#### Thermosynechococcus sp. Strain NK55a

had been sequenced recently and published 2014. An overall coverage of 26x was achieved with a final length of 2,519,964bp. This strain was isolated from the Nakabusa hot spring in Japan. It was cultured at 52-60°C. The 16s DNA sequencing results were almost identical to the *T. elongatus* strain.^54^

Significant cyanobacteria genomes belonging to the *Synechococcus* genus can be found in the following list. From today a further strain belonging to this small group can be added to this list.

**Table 2.** Significant cyanobacteria whole genome projects (Cyanobase/NCBI)

|  |  |
| --- | --- |
| *Thermosynechococcus elongatus* BP-1 | Kazusa (2002)" |
| *Thermosynechococcus* sp. **Strain NK55a** | Pacific Northwest National Laboratory (2014) |
| *Synechococcus* sp. WH8102 | JGI (2003) |
| *Synechococcus elongatus* PCC 6301 | Nagoya U. (2007) |
| *Synechococcus* sp. CC9311 | TIGR (2006) |
| *Synechococcus* sp. PCC 7002 | Penn State University (2008) |
| *Synechococcus* sp. CC9902 | DOE Joint Genome Institute (2007) |
| *Synechococcus* sp. CC9605 | DOE Joint Genome Institute (2007) |
| *Synechococcus elongatus* PCC 7942 | DOE Joint Genome Institute (2007) |
| *Synechococcus* sp. JA-2-3B'a(2-13) | TIGR (2007) |
| *Synechococcus* sp. JA-3-3Ab | TIGR (2007) |
| *Synechococcus* sp. RCC307 | Genoscope (2007) |
| *Synechococcus* sp. WH 7803 | Genoscope (2007) |

## Materials & Methods

In the workflow bellow the applied strategy for the whole genome sequencing project is visualized. The various stages required different techniques and methods, which will be introduced in the following chronological order.

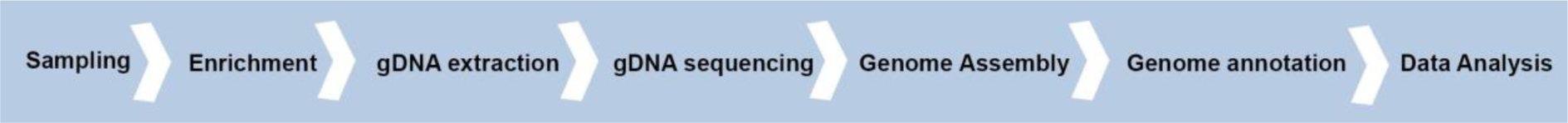

### Sampling of cyanobacteria

The sampling was undertaken in Namibia, (Southern Africa) on a farm called Klein Barmen at a hot spring nearby the city of Okahandjia. The farm owner/representative provided access to the fountain and this was confirmed to the local authority. The sample was taken at the GPS coordinates: S 22°08’’ 34.8’ E 14° 31’’ 33.0’ The temperature at the sampling site: 58°C. Fresh water with pH: 7

A small lake surrounded the fountain. Some flora were observed which are not typical seen in that environment. In figure 20 original photos of the sampling site are shown.

**Figure 20.**
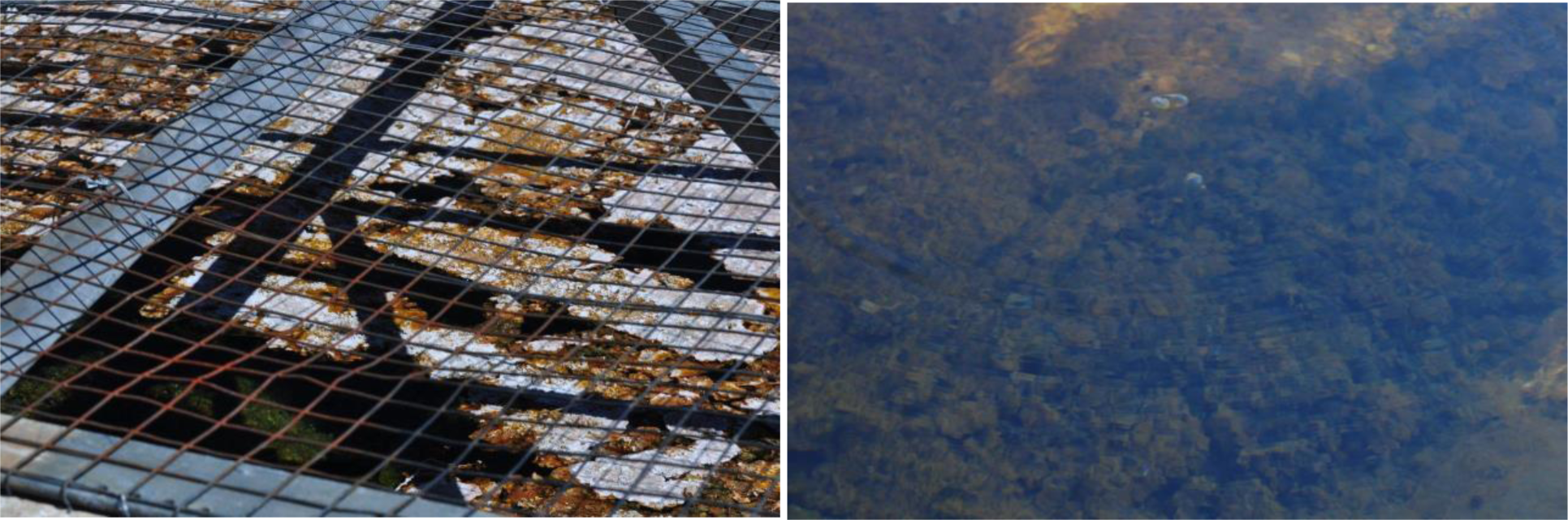
Cyanobacteria sampling site. Mats on surface (left) mud and debris at the bottom (right)

### Cultivation of cyanobacteria

In general many different protocols for cyanobacteria cultivation under laboratory conditions were established in the past. A wide range of different media formulations are described in literature.^45^ Most commonly cyanobacteria are cultured in cheap Erlenmeyer flasks under controlled artificial light. A current limitation to cyanobacteria research is that only a limited fraction of the natural diversity is cultivable under axenic conditions.^55^ A major requirement was to establish an axenic stable culture of the isolated thermophilic cyanobacteria.

### Cultivation of an environmental sample

In order to cultivate cyanobacteria present in the environmental sample various growth media were tested. 1 ml of the original sample was taken to inoculate 50 ml sterilized media in an Erlenmeyer flask. These flasks were then incubated for 2 weeks under low light. In the given table 3 the primary cultivation strategy is presented.

**Table 3.**
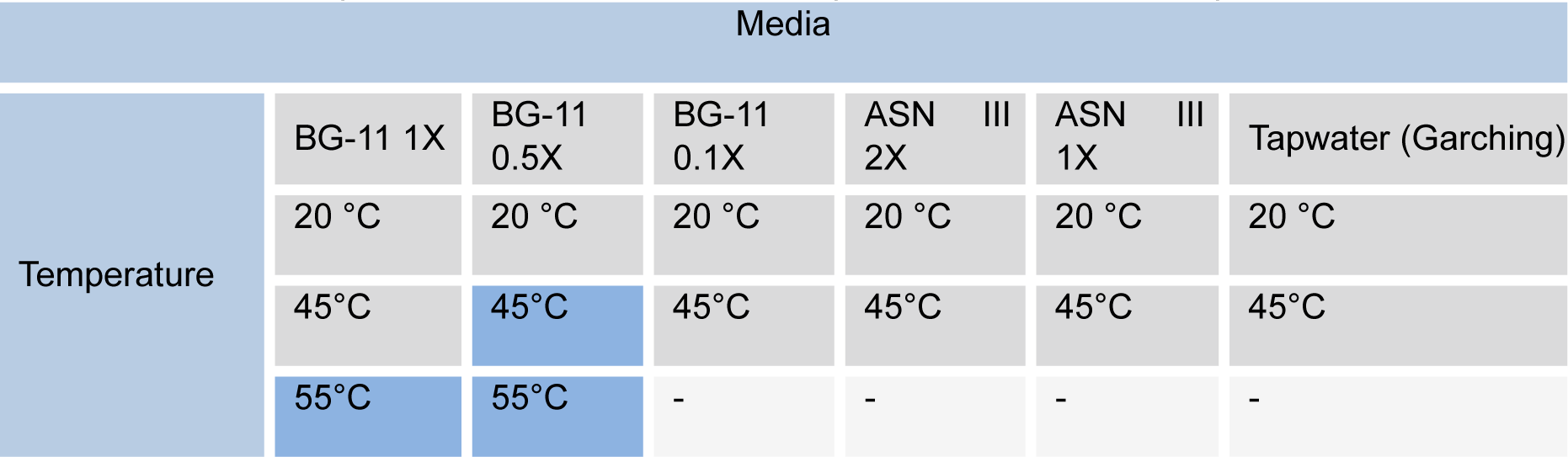
Tested culturing conditions. In blue the culturing media used for axenic growth.

### Establishing an axenic culture

For every whole genome sequencing project, enrichment is of importance to improve the signal noise ratio (SNR). Therefore, it was necessary to achieve pure culturing conditions. The standard method for cyanobacteria isolation is to perform dilution plating. Different solid media formulations were tested over the course of the project.

|  |  |
| --- | --- |
| **<u>Matrix builders:</u>** | Agar; Agarose; Washed Agarose; Washed Agar; Gelatine |
| **<u>Strength:</u>** | 0,4 -1,0% |
| **<u>Media and concentrations:</u>** | BG-11 0.5 x – 2 x |
| **<u>Special nutrients:</u>** | Thiosulfate; Vitamin B12; CaCO<sub>3</sub>; |
| **<u>Inoculum:</u>** | 1 ml – 10 µl by plating / streaking / spray inoculation |
| **<u>Carriers:</u>** | Petri dish (Sarstedt small 7cm polystyrol / glass); Erlenmeyer flasks |
| **<u>Temperature:</u>** | 20°C / 50°C |

Dilution plating was not successful for isolation, therefore another approach had to be considered. With a Biorad FACS device, single cell sorting was used to obtain axenic cultures. Cyanobacterial photosystems (chl a) absorb light with discrete peaks around 700 nm (+/− 70) ( Biorad filter configuration). This was used to sort efficiently chlorophyll containing cells from “contaminants”. In order to sort the cells, an appropriate dilution of a fresh culture was prepared to meet the sorters specifications. The sorter was calibrated for single cell sorting. 2 ml Eppendorf tubes filled with 1.5 ml BG-11 media were used as “receivers”. The Eppendorf tubes were incubated thereafter in a rack under 20 μmol m^−2^s^−1^ constant white (4000K) LED radiation. The tubes were closed and the underneath residual radiation was measured. After 1 week of incubation the cultures were passaged into larger culturing vessels. In the following diagram the workflow is depicted.

**Figure 21.**
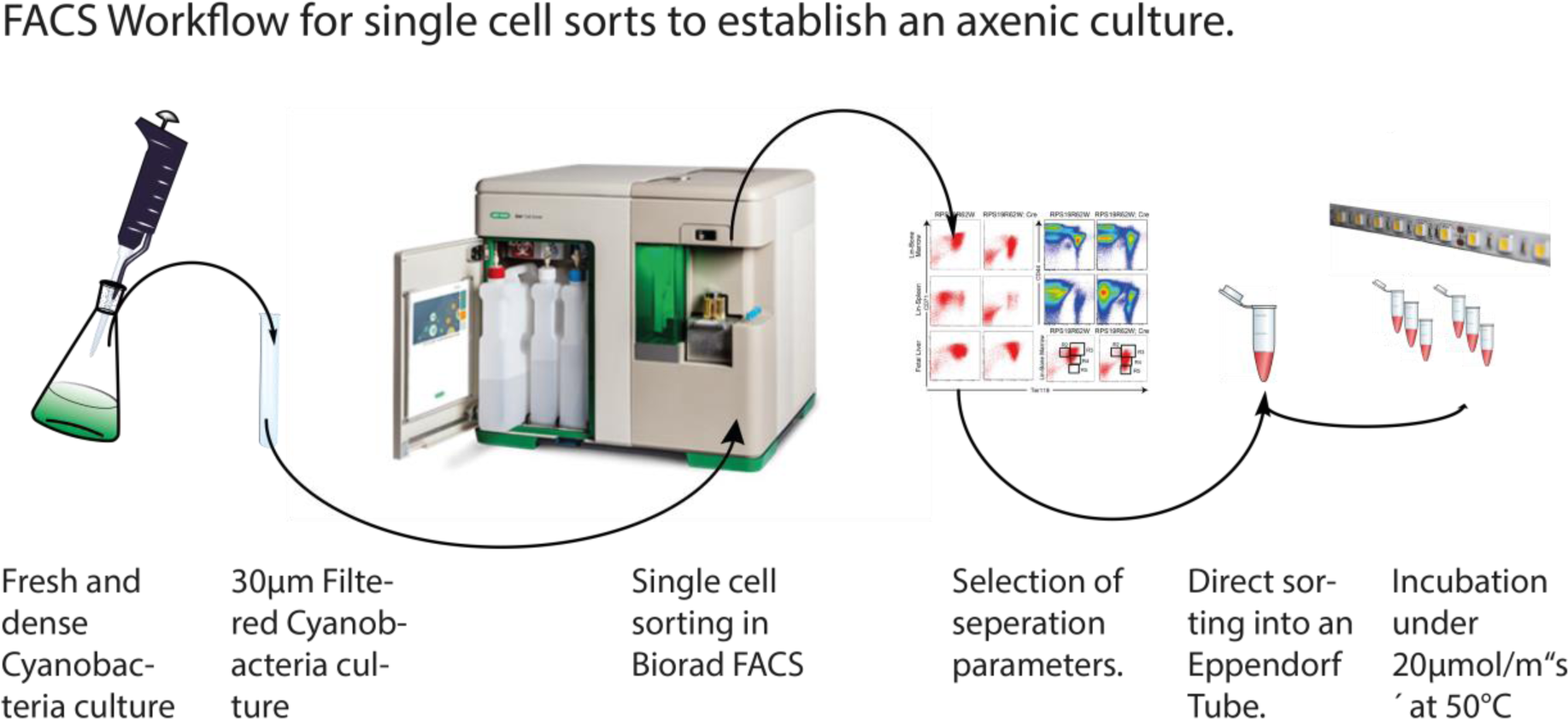
The FACS workflow. The sample was filtered with a sterile cell strainer cap. Thereafter, diluted in sterile media and placed in the Biorad FACS device. The sorting gates were applied as described in the results chapter.

As cyanobacteria are relatively small and have a gram-negative stable cell wall, cyroconservation was possible. The cryoconserved samples of pure isolates were stored at - 80°C and used throughout this project.

### sDNA sequencing for phylogenetic classification

By comparing literature morphologies it is possible to categorize sample composition by classical microscopic means. A much greater depth of characterisation can be easily achieved by 16s (or 18s for Eukaryotes) rRNA comparison by NCBI databank BLAST^56^. This method has revolutionized microbiology in terms of taxonomic classification. For all 16s rRNA BLASTs^56^ we used NCBI – to include all possible genomic sequences related to this cyanobacteria.

Primers which are known to be consistent over all bacteria species were used^57^ Primer forward: 27f AGA GTT TGA TCA TGG CTC A - l: 19 Tm: 52.4 GC 42 Primer reverse: 1492r TAC GGT TAC CTT GTT ACG ACT T - l: 22 Tm: 56.5 GC 41 With this primers PCRs of the enriched sample were conducted. For this 25 ml of a 1 week old culture was centrifuged with ∼ 10 000 rpm. The supernatant was disposed the pellet re-suspended in 50 µl media and incubated for 10 min at 95°C. The lysate was thereafter used as input for the PCR. The PCR was run by a standard amplification protocol using an annealing temperature of 62°C.

The PCR product was analysed and purified by gel electrophoresis. The gels were run with standard 100 Volt and 5 Watt for 30 min in Biorad gel cambers filled with TAE buffer. After gel electrophoresis the appropriate bands were cut, digested, purified and concentrated using Analytic Jena InnuPrep kit. After a photometric quality control 15 µl sample (min 10 ng/µl) with 1 µl primer (27f and 1492r) was sent to the local sequencing service.

The results of the Sanger sequencing were BLAST^56^ searched against the NCBI database; using BLASTN 2.2.32+ ^56^ against all nucleotide entries.

After DNA sequencing and assembling the *de novo* genome, a further more accurate 16s rRNA study was performed with the MinION™ data. The obtained 16s rRNA ABI sequence was mapped by Geneious against the MinION™ *de novo* genome. The overlap region was then used for another NCBI BLAST query.

### Oxford Nanopore MinION™ d*e novo* genome sequencing

During this study the novel nanopore DNA sequencing device was used to sequence genomic DNA. In order to conduct NGS sequencing, high quality DNA has to be extracted. Once the DNA extraction is conducted, a DNA library to be loaded “on” the sequencing technology has to be prepared. The successfully prepared library is then “loaded” on to the device and the data was further processed after completing the sequencing run.

For genomic DNA (gDNA) extraction, an axenic dense culture was cultured to stationary phase. The genomic DNA extraction was done with the ONT recommended Qiagen genomic DNA extraction kit (QIAGEN Genomic-tip 500/G cat: 10262). The protocol for bacterial DNA isolation was performed. The sample input was calculated by a cell-count / OD600 correlation, to minimise the risk of overloading the column. A few adaptions to the protocol were made: incubation time for the lysis of the bacterial cells was increased to 75 min. The lysate was vortexed for 30 s. After elution from the Qiagen Genomic-tip 500 (G-tip) the extract was precipitated twice with EtOH and dissolved in 1 ml ddH2O for 3h at RT on a shaker.

After the genomic DNA extraction with Qiagen G-tip, the quality of the isolated DNA was verified by photometric Nanodrop™ and flourometric Qubit™ measurements. The extracted DNA was further purified by EtOH precipitation. Finally, the concentrate was cleaned from residual impurities with the Analytic Jena purification kit, (innuPREP Gel Extraction Kit 845-KS-5030250) resulting in high quality ultra-pure gDNA.

The gDNA input for all sequencing runs was taken from the same extract over a course of 2 weeks. The excess gDNA extract has been preserved for future re-sequencing projects.

Generally, there are various DNA extraction methods to choose from. This should be kept in mind if the input DNA quality is not satisfactory. Nanopore sequencing requires highly pure samples. The following table lists an overview of DNA extraction methods. In grayscale are the purification methods used during this work.

**Table 4.** DNA purification methods translated from der “Experimentator Genomics”^58^.

| DNA purification methods |  |
| --- | --- |
| PEG- precipitation | Phenol–chloroform extraction |
| Anion-exchange column | Dialysis |
| Magnetic beads | Silica membrane |
| Ethanol- precipitation | Isopycnic centrifugation |
| Filtration (Amicon) | Protein binding filter membrane |
| Butanol /extraction | Solvent extraction |

After DNA extraction and purification, the library preparation for nanopore sequencing was conducted. For the library preparation the developed checklist type ONT protocol was used. In the following the most important steps are presented. No alterations to the protocol were made.

To ease up bioinformatics bottom up analysis, and the sequencing process itself, the DNA library has to fulfil some basic criteria. The length distribution of the DNA fragments has to be as sharp as possible. More important, the library has to be pure and free from undesired DNA contaminations which interfere with the sequencing technology.

The Qiagen^TM^ raw extract has a length distribution of about 100-150 kb.^59^ This can be assessed with Agilent’s Bioanalyser for trouble shooting and quality control (QC) if available.^1^ For the MinION™ a size distribution around 8 kb - 10 kb is producing currently the most reliable and high throughput results.^20^ Therefore, gDNA is sheared in a reproducible manner to produce normal disturbed fragments. To shear the gDNA to 8 kb fragments the recommended Covaris™ g-tubes were used. These tubes allow the fragmentation of the extracted DNA by centrifugation. Different length distributions can be created by varying the centrifugation speed. The consumable is essentially a plastic tube with a small filter/frit in the centre. By centrifugation the DNA solution is forced through the filter/frit and sheared by it subsequently.

Important is that no PCR step was necessary before – a third generation sequencing characteristic.^14^ After Covaris™ shearing, the library was end repaired using the NEBnext end repair kit.

The enzyme/DNA mixture was purified with Beckmann coulters Ampure™ beads by magnetic sedimentation. After elution from the Ampure™ beads, the library was dA tailed with the NEBnext dA tailing kit. The enzyme mix ligates a dAMP at the 3’ blunt end. A further Ampure magnetic bead purification step was done. The dA tailed library was then ligated to ONTs special adapters “AMX/HPA” needed for the strand progression through the nanopore. To stay in the recommended concentration of 0.10.2 pmol of gDNA free ends, the initial input mass of gDNA was adapted according to the Covaris g-tube fragmentation length. After ligation of the DNA fragments containing the motor enzyme, care had to be taken during handling not to denaturate the sensitive complex. Therefore, pipetting steps were minimised and protein low bind Eppendorf tubes were used. The library was finally purified by “Histag” Dynabeads. The “dynabeads” are binding ligation products with hairpin adapters (HPA have a protein bound with terminal histidines). After elution of the purified library from the Histag beads, the so-called pre sequencing mix (PSM) was stored on ice/4°C. A final quality control step was done with a qubit fluorimeter (life technologies) determining PSM library (DNA) concentrations most accurately.

Before loading the library on the MinION™ the mobile workstation had to be prepared for a 48h continuous operation. (This cannot be overemphasised as it did trick some fellow mappers.) After checking for software updates – the software MinKNOW which controls the MinION™ device had to be launched as well as the Metrichor cloud based base calling service. Both were working in parallel and in real time. After mounting the device on the PC a quality control of the flow cell had to be conducted. This step assesses the amount of intact protein pores able to analyse DNA strand progression. After quality control the flow cell needed to be primed with a running buffer and fuel mix (FMX). The priming step was done with 300 µl “running buffer” before loading of the sequencing mix (S.M.) on the flow cell. The final S.M. was then loaded on the flow cell on the MinION™ device by a 1000 µl pipette. Directly after loading of the S.M. the appropriate python script was started. Every six hours new S.M. was loaded; until no PSM (Pre-sequencing mix) was left.

Instantly the sequencing signals are saved and directly transferred to the Metrichor™ base calling service. After basecalling the files are downloaded into a local destination directory. The files are sorted into either a “pass” file (PF) or a “fail” file (FF). The pass folder does include all data which were high quality 2D reads, generated by the MinION™ and Metrichor basecalling algorithm. The runtimes varied from 24-48 h depending on the flowcell performance (or required data).

In the following “workflow diagram” in figure 22 the library preparation steps are described in detail. The handy laboratory checklist protocol chart were created from the lengthy protocol over the course of this study. It includes all relevant information for the conduction of the library and serves as documentation for every run. Different versions of this document are updated by ONT regularly as a tool for quality control. I highly encourage using this tool to increase reproducibly and comparability between different laboratories.

**Figure 22:**
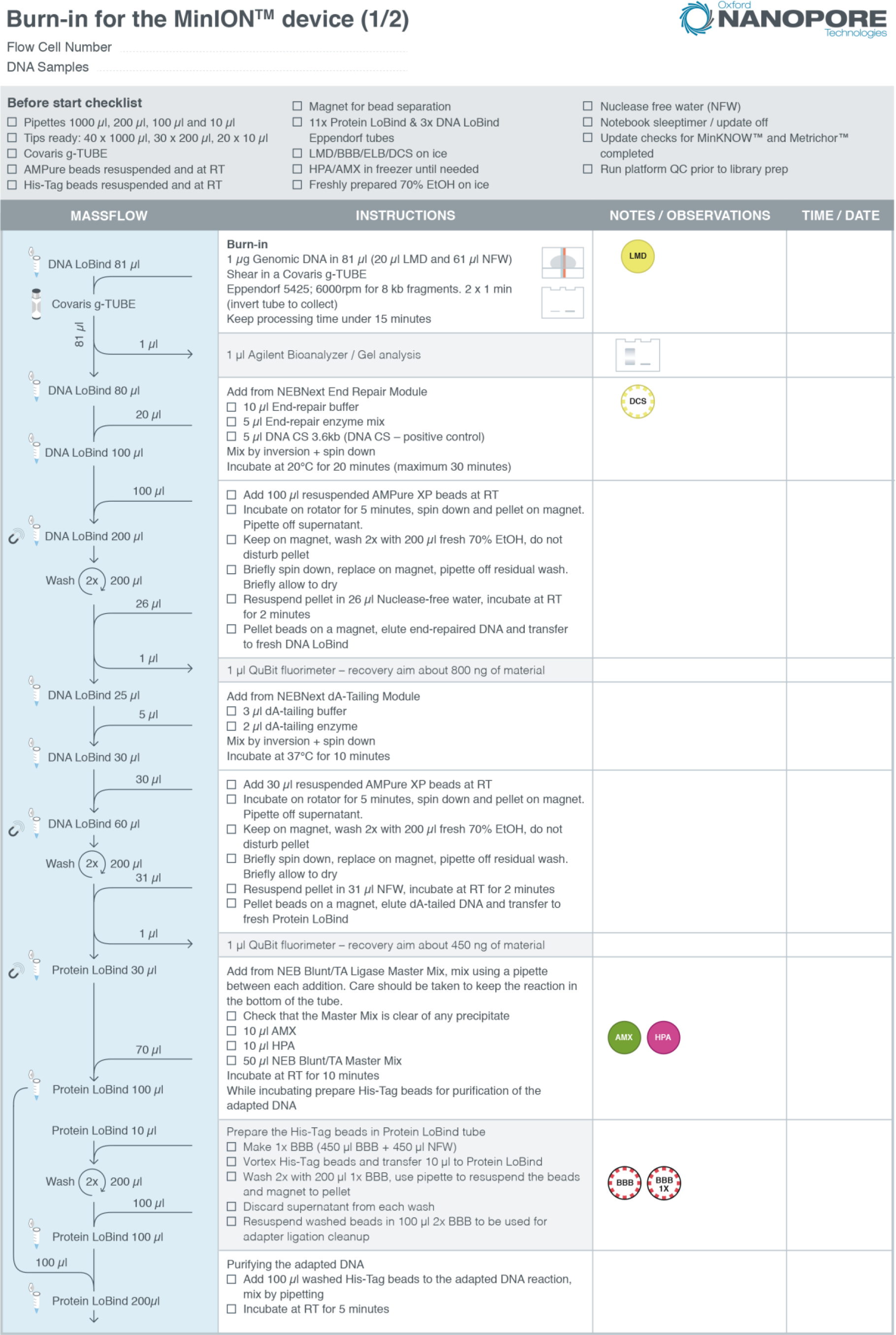
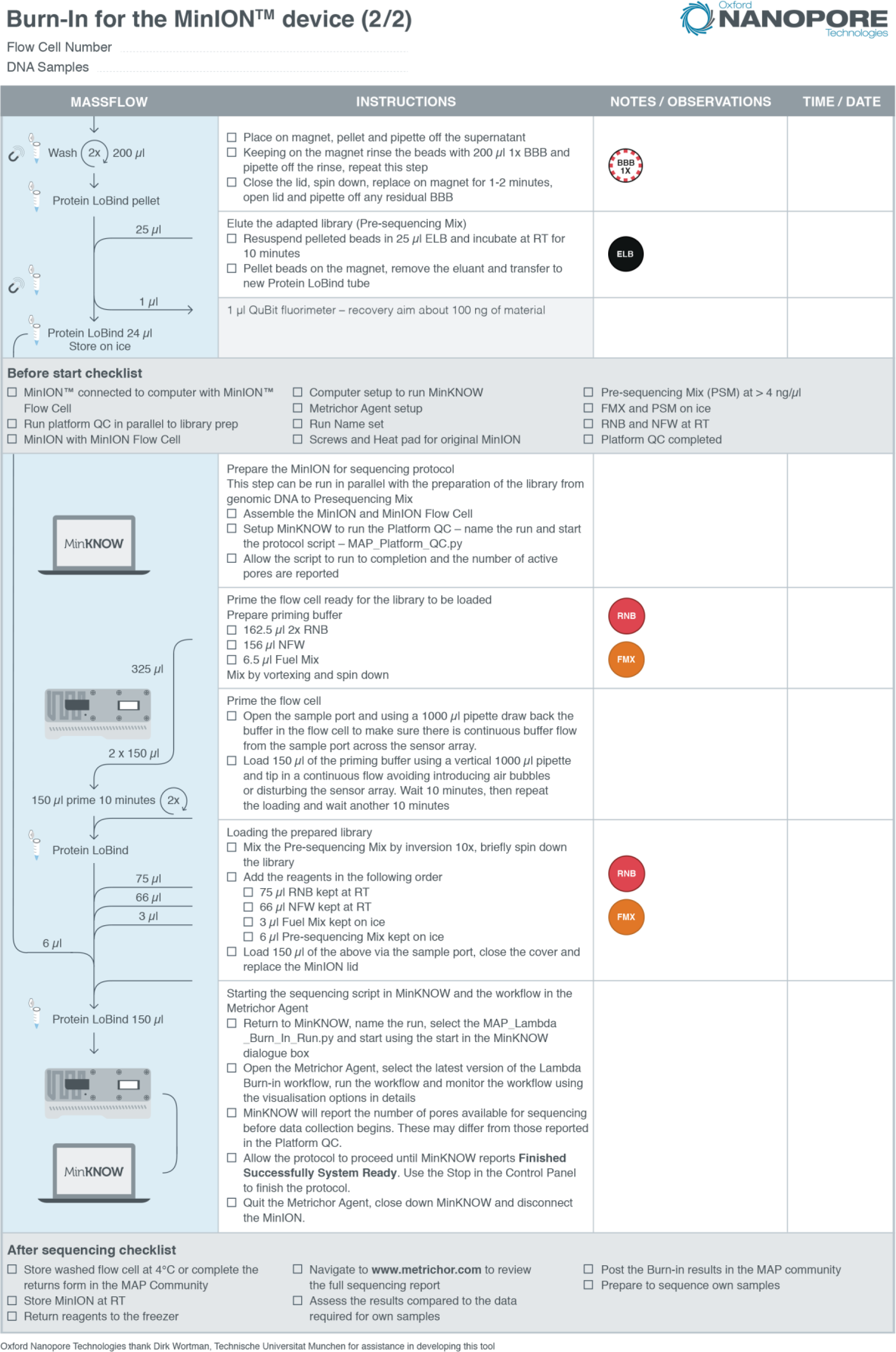
Workflow checklist for R7.3 Chemistry 08/2015. The tool was developed over the course of this study and provided to Oxford Nanopore Technologies™.

### Assembling the Synechococcus genome

After successfully sequencing with the MinION™ the data was used to perform a *de novo* genome assembly. The computational work was done with the help of the IBK server computer and a Lenovo mobile workstation. On both computers virtualized Ubuntu 14.04 on Win 7 Pro was used.

For this task various software packages were installed and run. In figure 23 at the bottom the different workflows and single software tools are documented.

**Figure 23.**
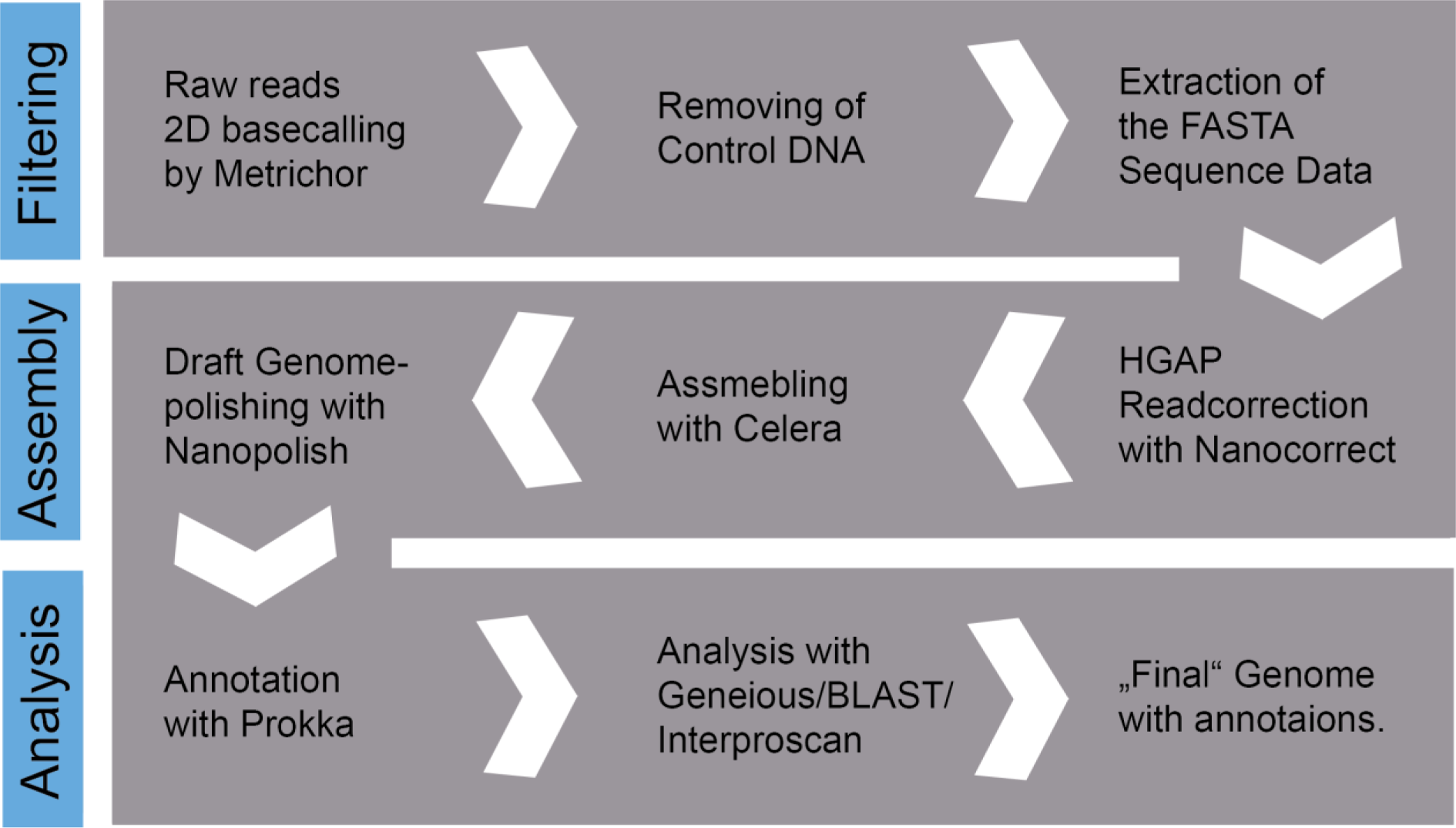
The NGS analysis Workflow. Read extraction was done with Poretools^60^.

The raw data downloaded from Metrichor passing 2D basecalling and quality control, were stored in a folder containing all.fast5/HDF type files.

A further data filtering step was used to remove control DNA reads. This was done by Mirabait^61^, finding all reads homologous to the control DNA. The corresponding reads were excluded thereafter. To data which were filtered by this method a CF prefix was added.

Thereafter, the fast5 files were processed with Poretools^60^. Poretools is an R^62^ package freely available enabling FASTA and FASTQ extraction from FAST5 as well as some analytical tools to evaluate the sequencing run success.

The extracted FASTA data was then corrected using Nanocorrect^2^ – a toolkit from Loman et al. Nanocorrect is a software tool compromising an alignment and correcting approach in accordance with the HAGP.

After this correction pipeline the data was assembled using Celera^19^. After assembly with Celera the draft genome was polished using the Nanopolish^2^ toolkit. Nanopolish does as the name infers, correct the assembled genome once again. Nanopolish uses a HMM (Hidden Markov Model) to correct the data according to the raw signal data included in the FAST5 files. The resulting “polished” genome was then analysed and annotated.

### Annotation of the *Thermosynechococcus DVW* genome

Genome annotation was defined by Richardson and Watson in the following way: *“Genome annotation is the process of identifying and labelling all the relevant features on a genome sequence”* ^63^

In order to assign functions to the newly gained genome, annotation software was used to predict possible proteins and regulatory elements. The *de novo* genome was annotated by Prokka^64^ – an automated annotation software tool for bacterial genomes (Prokka^64^ Torsten Seemann / Monash Bioinformatics Platform). The tool was run on a mobile workstation. The input data requirement is the polished FASTA file only. The output of the annotation process was directly edited and analysed in Geneious software suite. The Prokka^64^ software uses 5 different tools and services to annotate the sequence. These are listed below:

**Table 5.** List of tools ^64^ implemented in Prokka ^64^.

| Tool (reference) | Features predicted |
| --- | --- |
| Prodigal (Hyatt 2010) | Coding sequence (CDS) |
| RNAmmer (Lagesen et al., 2007) | Ribosomal RNA genes (rRNA) |
| Aragorn (Laslett and Canback, 2004) | Transfer RNA genes |
| SignalP (Petersen et al., 2011) | Signal leader peptides Non-coding RNA |
| Infernal (Kolbe and Eddy, 2011) | Non-coding RNA |

## Results

An overview of all significant results are presented and listed in the following chapter. This will be the basis for the discussion focusing on MinION™ sequencing data and the novel thermophilic cyanobacteria genome.

### Isolation and cultivation of a wt cyanobacteria

The environmental sample was cultured as described in 4.2.1. Growth of the sample was observed only in 0.5x BG11 at 40°C. The culture was passaged every two weeks into fresh media to maintain stable growth. At unfavourable conditions and in old cultures reaching stationary phase, it was often observed that the cyanobacteria turned to a dormant stage. As a routine cell morphology and purity control method, light microscopy was adopted. In the following diagram examples of some microscopy images from the studied strain are presented. The cyanobacteria is approximately 4,7 µm long and 1.5 µm in diameter.

**Figure 24.**
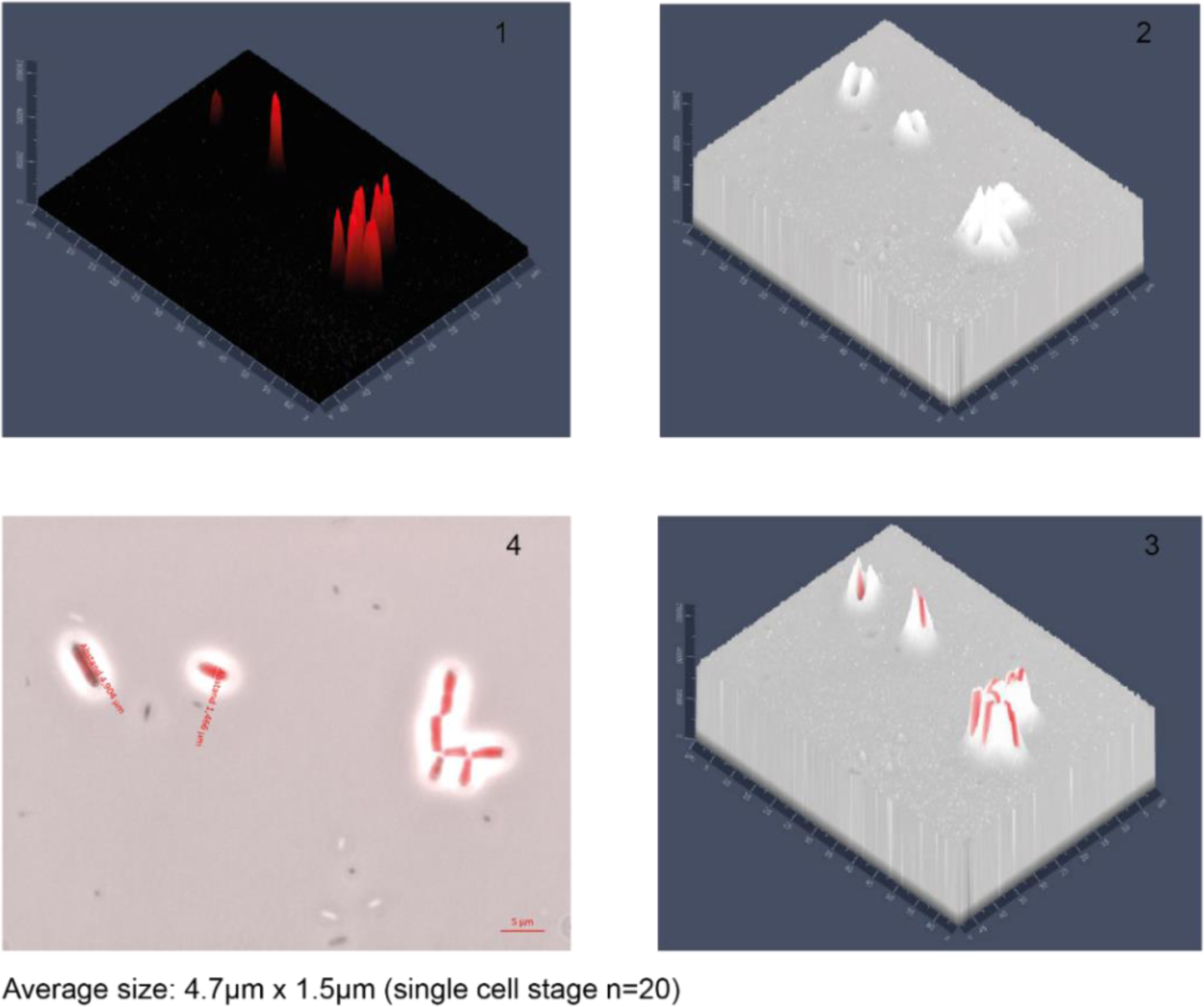
P2NDW WT Cyanobacteria 1000x (Zeiss Axio Lab); 1: 3D Chlorophyll LP filter (excitation: 480/36 nm. emission: 635 nm); 2: 3D Bright-field PH3 Picture 3:3D Chl. / Bright field PH3; 4: 2D Chl. / Bright field PH3 Cultivation and establishing axenic growth

Establishing axenic culture is still a requirement for *de novo* genome assembly by MinION™ sequencing. Therefore, different approaches have been carried out to gain a pure “homogeneous” culture. Cultivation on solid media was not successful, although various conditions were tested. To overcome this limitation, the widely used FACS method for single cell characterisation and counting was used. In the following figure 25 the “gates” for cell sorting experiments which yielded pure culture isolates are shown.

**Figure 25.**
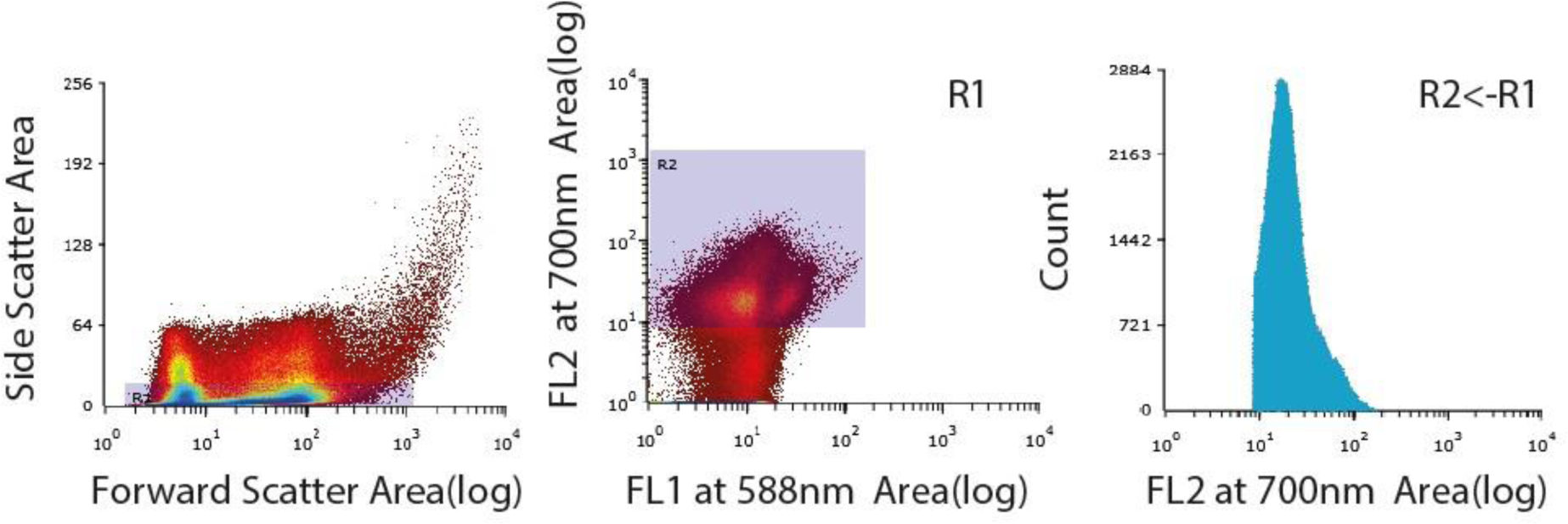
FACS Cyanobacteria isolation settings. Left: The sorting gate side scatter was applied to exclude debris. The sorting gate shown in the centre selected high FL2 candidates.

The single sorted cells were cultured in 1.5 ml Eppi and after a week passaged into an Erlenmeyer flask with 25 ml BG11. The negative control did not show any growth. The fastest growing samples were selected and cultured. Two cultures (No.13 and No. 27) showed the best stable long-term growth and were considered for the NGS experiments.

### s rRNA sequencing

In order to verify and define the taxonomy a 16s rRNA sequencing study on the enriched culture was conducted. The corresponding clipped 16s rRNA sequence were “blasted” against all NCBI nucleotide collection (nr/nt) entries. The 16S rRNA (DNA) sequence can be found in appendix 10.4. The BLASTN 2.2 ^56^ alignment results of “the sequence are presented in the table 6.

**Table 6.** Results Megablast^56^ Query_1 of the PCR Sequence.

| Description | Total score | Query cover | Ident. | Accession |
| --- | --- | --- | --- | --- |
| Uncultured *Synechococcus* sp. clone YCB100 | 1463 | 99% | 99% | EF205441.1 |
| Uncultured bacterium gene for 16S rRNA, | 1458 | 99% | 99% | AB757741.1 |
| *Synechococcus* sp. CR_L35 16s | 1426 | 97% | 99% | EF545646.1 |

After *de novo* assembly of the genome the corresponding 16s rRNA sequence was used to run a second NCBI BLAST search. The results are presented below in the table 7.

**Table 7.** Results of Megablast^56^ Query_119085 with 16s rRNA sequence extracted from the assembled genome.

| Description | Total score | Query cover | Ident. | Accession |
| --- | --- | --- | --- | --- |
| *Synechococcus lividus* C1 | 1441 | 100% | 98% | AF132772.1 |
| *Synechococcus* PCC6716 | 1441 | 100% | 98% | AF216942.1 |
| *Synechococcus* PCC6717 | 1435 | 100% | 98% | AF216943.1 |
| *Candidatus Synechococcus calcipolaris* | 1413 | 100% | 98% | KJ566932.1 |
| *Thermosynechococcus* sp. NK55 genome | 1402 | 100% | 98% | CP006735.1 |
| *Thermosynechococcus elongatus* WFW | 1397 | 100% | 97% | KC621876.1 |
| *Thermosynechococcus elongatus* BP-1 | 1397 | 100% | 97% | NR_074328.1 |
| *Synechococcus elongatus* | 1397 | 100% | 97% | D83715.1 |

The Megablast^TM^ results from table 7 were visualised using the NCBI phylogenetic tree function. The tree parameter was set to: fast minimum evolution, max seq. difference 0.05.

**Figure 26.**
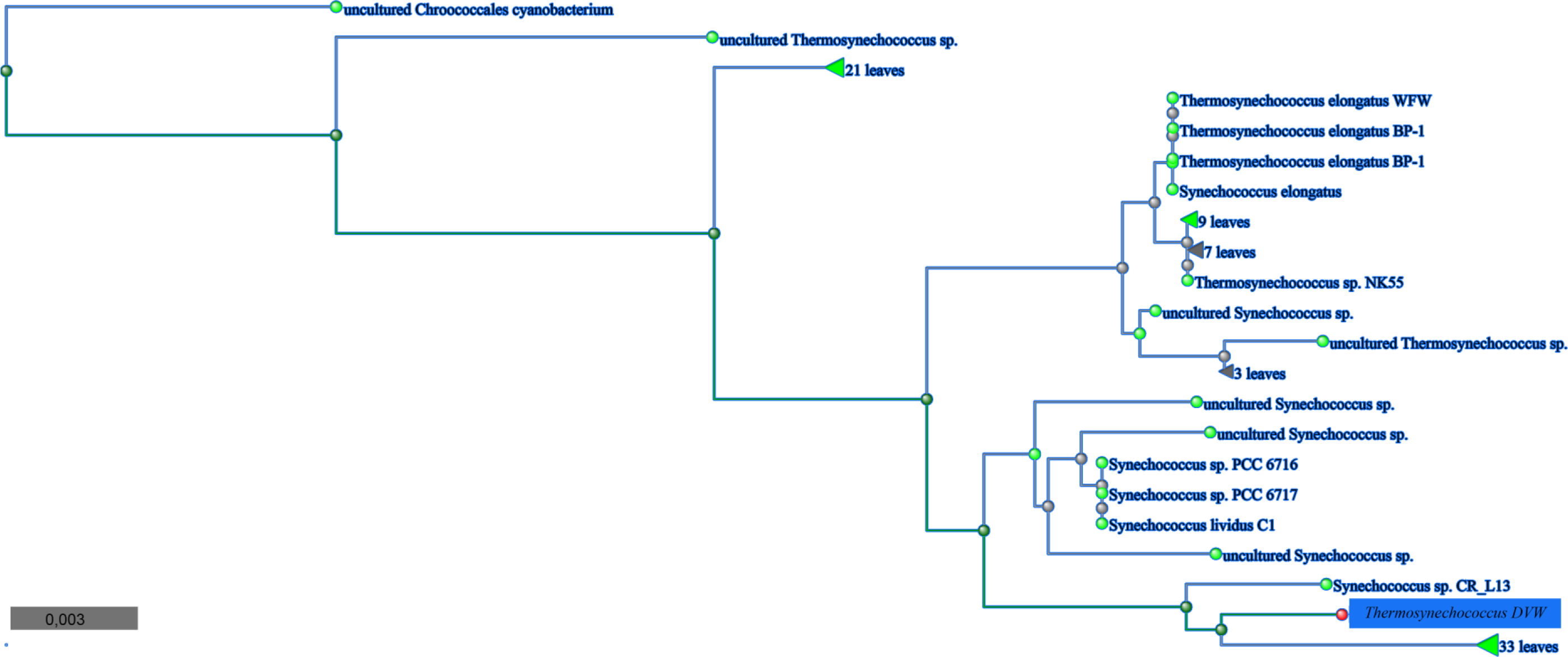
NCBI Phylogenetic tree, Megablast^56^ results for 16s rRNA Sequence obtained from the de novo genome. (Modified by Adobe Illustrator.

### Genomic DNA isolation and purification

In order to estimate the cell-count (gDNA copy number) an OD600 cell number correlation was determined with “Improved Neubauer Chamber” cell counting.

**Figure 27.**
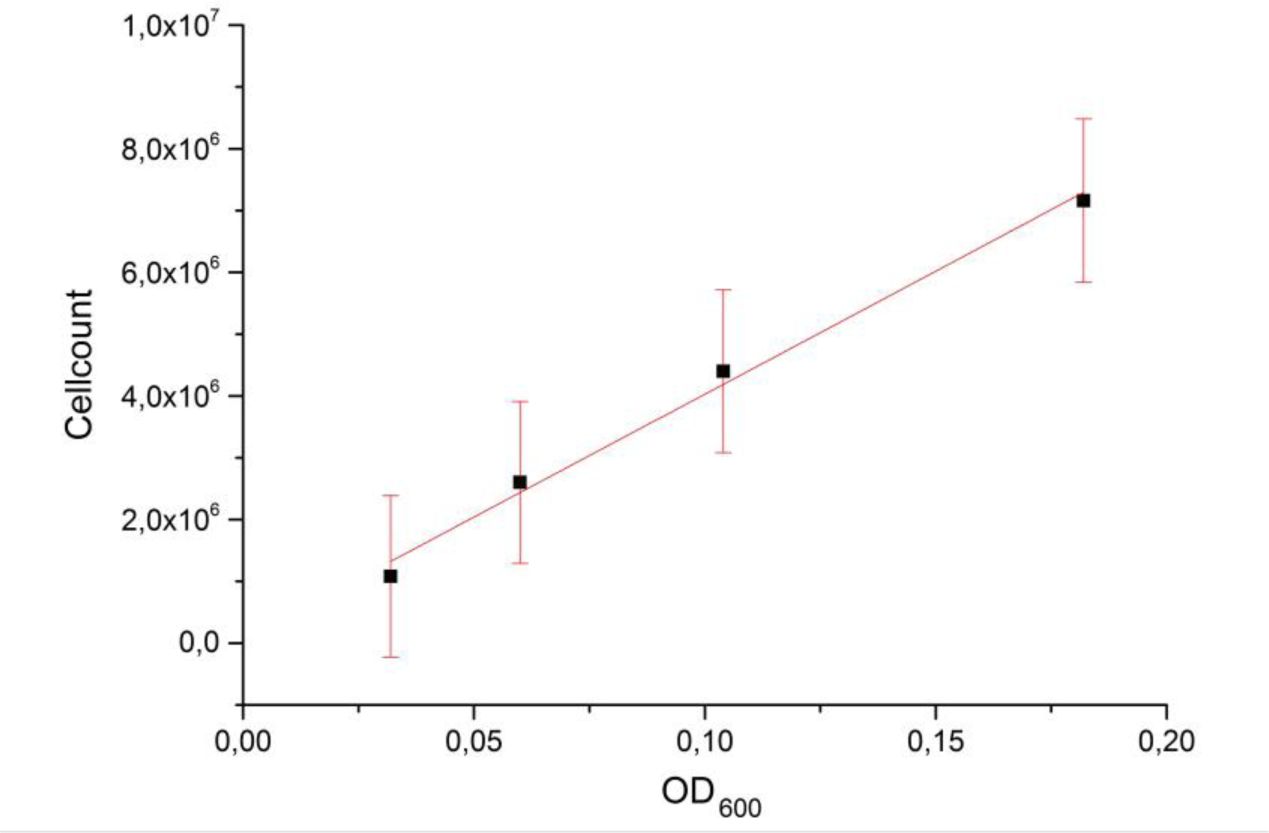
OD600 cell-count correlation to estimate the culture volume needed for DNA extraction. The graph shows the linear dependency of cell-count and OD600 for Thermosynechococcus DVW. Plotted with Origin Pro.

From the cell-count / ml the necessary column capacity was estimated.

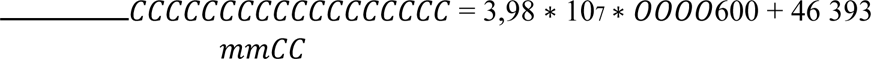

With the parameters from above a yield of 100 µg DNA extract was expected. For the extraction, 175 ml stationary sterile culture of No. 27 by a Qiagen G-tip 500 was used. Thereafter, EtOH precipitation and final InnuPrep™ purification followed.

The extracted gDNA quality was assessed and the final results are shown in figure 28.

**Figure 28.**
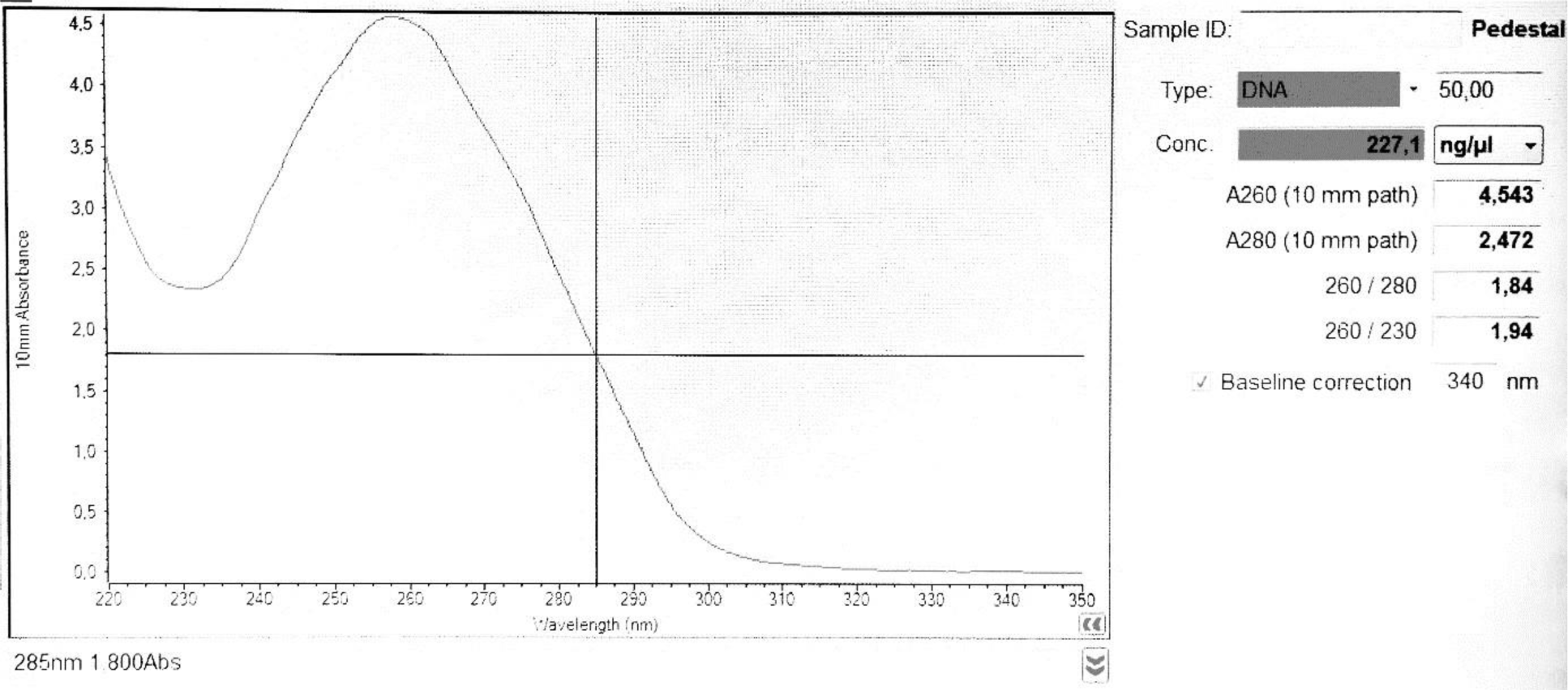
Nanodrop QC report for the genomic DNA used for the MinION™ runs.

To verify the fragment length of gDNA a low percentage agarose gel shown in figure 30 was used. ONT LMD DNA was used as control. The extract was stored until the library preparation for the MinION™ runs were conducted.

**Figure 30.**
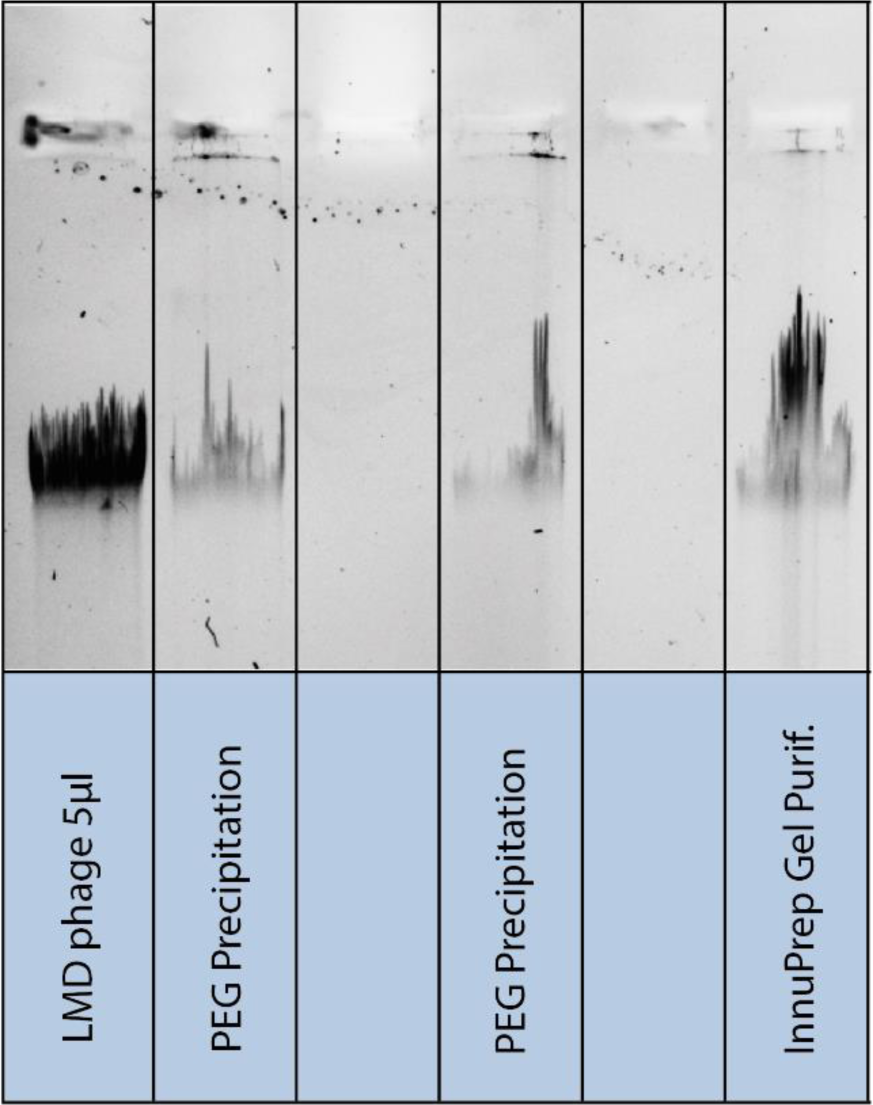
Low % Agarose Gel for gDNA QC. LMD phage DNA is the positive control at 48kb. The DNA corresponding to the InnuPrep lane on right was used as final Library preparation input.

### 5.4 Genomic DNA sequencing with the MinION™ device

After the DNA extraction as described in chapter 4.4, the genomic DNA was prepared for direct sequencing without PCR amplification. Three independent library preparations and MinION™ runs were conducted.

The following part of this chapter presents the data output and pre assembly processing results of the three independent runs. Every run which contributed to the assembly and analysis is included. In figure 31 the different data subsets and preprocessing results are visualized.

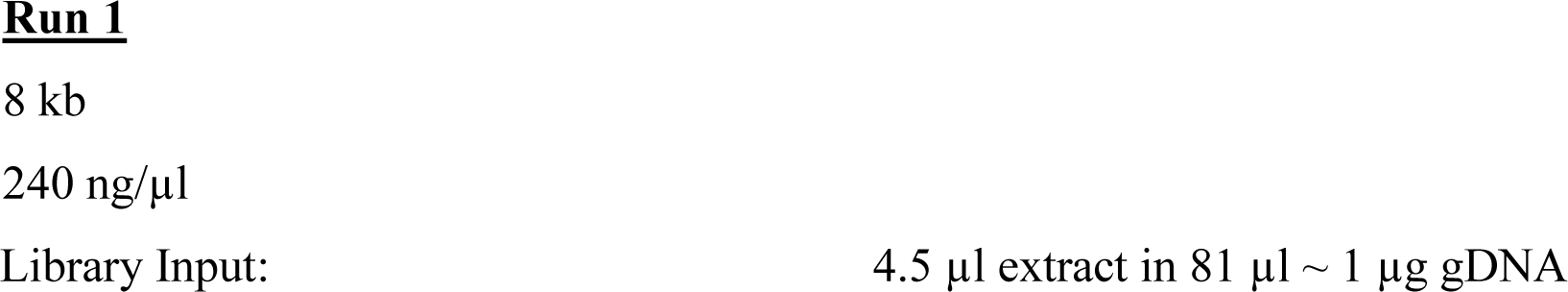

**Table 8.**
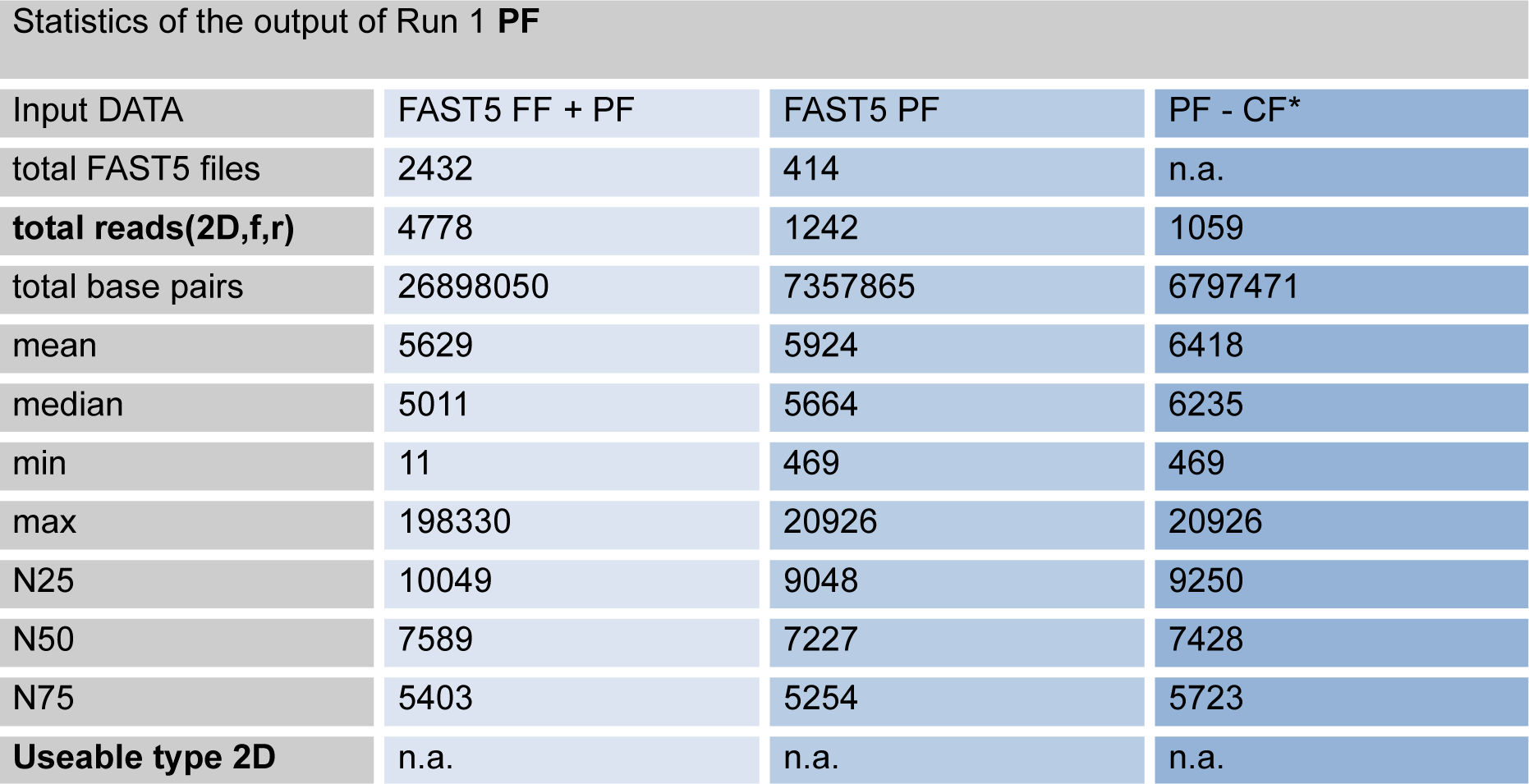
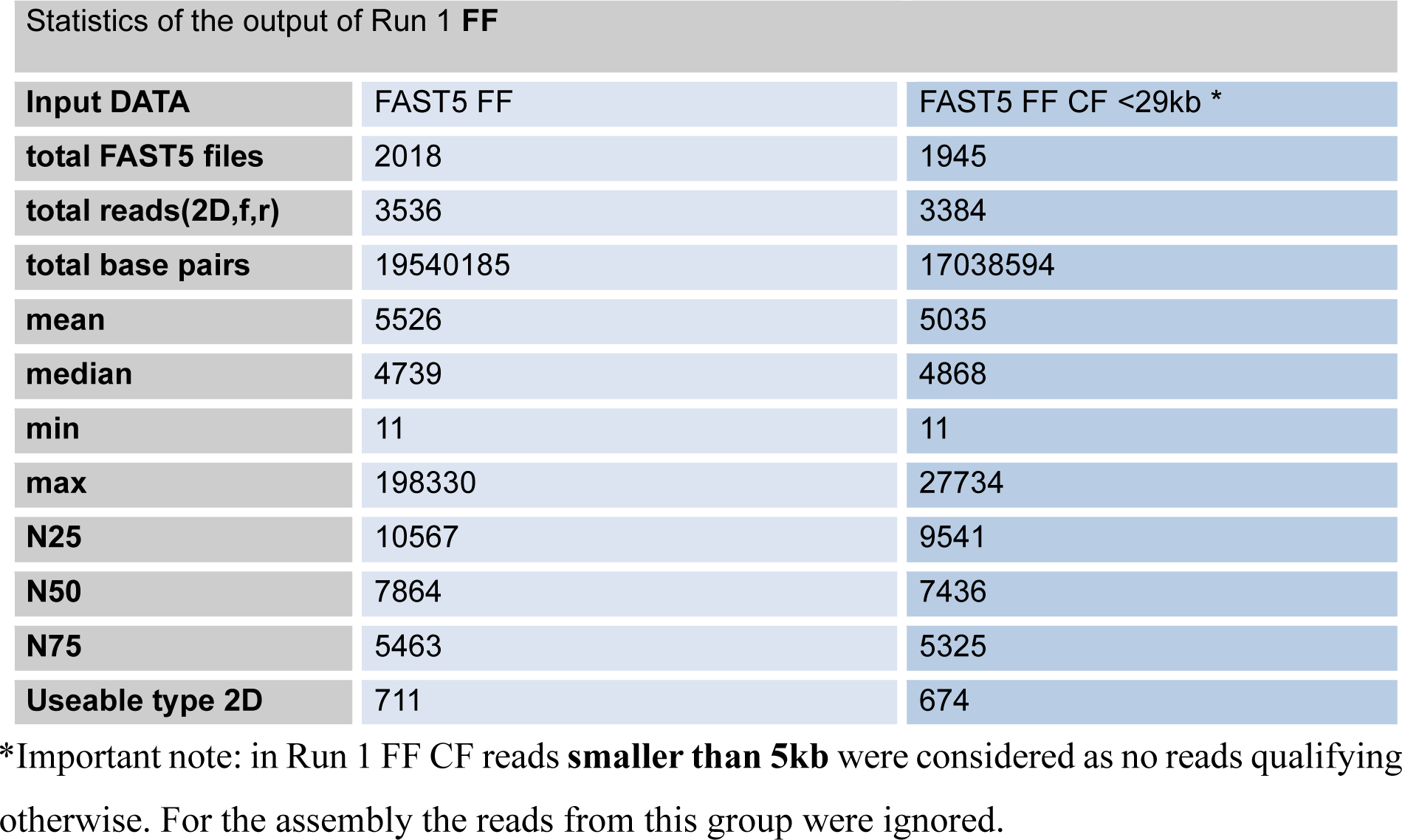
Run1 data from R *poretools*^60^ stats; CF = “control free”; PF = “pass filter/folder” (Metrichor) FF = Fail filter/folder (Metrichor); 2D = consensus from complement/reverse and template/forward sequenced strand (see introduction 3.2.2); f = forward; r = reverse

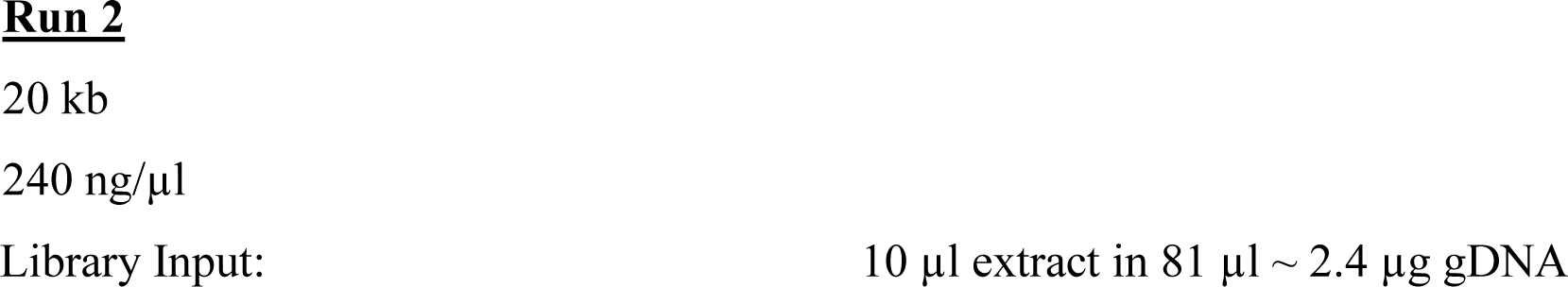

The library was eluted and QC resulted in 41 ng/µl PSM concentration. Total mass 1066 ng. 6 µl were used as input for the SM. Flowcell QC: 479 active pores

**Table 9.** Run 2 data from R *poretools*^60^ stats; CF = “control free”; PF = “pass filter/folder” (Metrichor) FF = Fail filter/folder (Metrichor); 2D = consensus from complement/reverse and template/forward sequenced strand (see introduction 3.2.2); f = forward; r = reverse

| Input DATA | FAST5 FF + PF | FAST5 PF | PF - CF* |
| --- | --- | --- | --- |
| total FAST5 files | 24463 | 3312 | n.a. |
| total reads | 41802 | 9936 | 1776 |
| total base pairs | 161264218 | 39691659 | 14179941 |
| mean | 3857.81 | 3994.73 | 7984.20 |
| median | 3169 | 3385 | 7984 |
| min | 6 | 157 | 157 |
| max | 296901 | 30742 | 30742 |
| N25 | 10400 | 7960 | 12011 |
| N50 | 5370 | 3475 | 9679 |
| N75 | 3268 | 3326 | 7549 |
| Useable type 2D | n.a. | n.a. | n.a. |

Statistics of the output of Run 2 **FF**
| Input DATA | FAST5 FF | FAST5 FF CF >5kb < 29kb |
| --- | --- | --- |
| total FAST5 files | 21151 | 4103 |
| total reads(2D,f,r) | 31866 | 6158 |
| total base pairs | 121572559 | 58227189 |
| mean | 3815 | 9455 |
| median | 2864 | 8587 |
| min | 6 | 5002 |
| max | 296901 | 28953 |
| N25 | 11116 | 12820 |
| N50 | 6438 | 9843 |
| N75 | 3198 | 7623 |
| Useable type 2D | 4465 | 1187 |

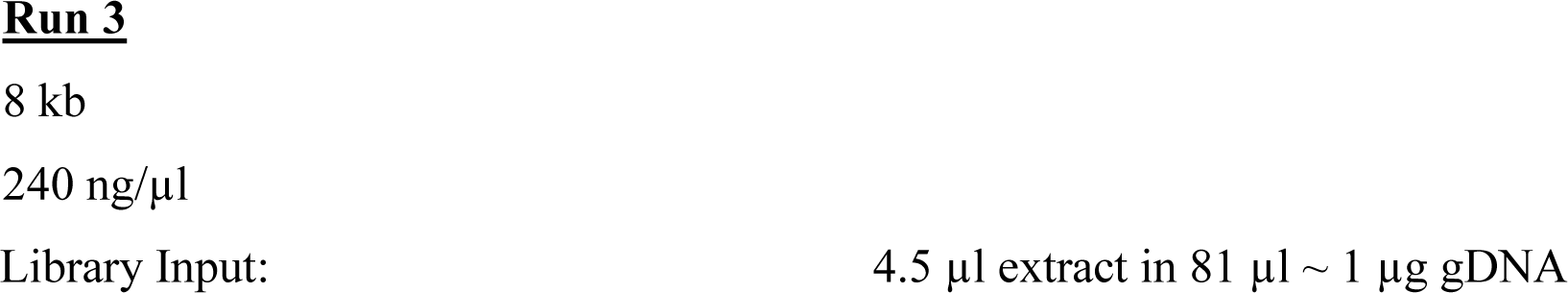

The library was eluted from the His-tag Beads and QC resulted in 9,6 ng/µl PSM concentration. Total mass 250 ng. 6 µl were used as input for the SM. Flowcell QC: 692 active pores

**Table 10.** Run 3 data from R *poretools*^60^ stats; CF = “control free”; PF = “pass filter/folder” (Metrichor) FF = Fail filter/folder (Metrichor); 2D = consensus from complement/reverse and template/forward sequenced strand (see introduction 3.2.2); f = forward; r = reverse

| Input DATA | FAST5 FF + PF | FAST5 PF | PF - CF* |
| --- | --- | --- | --- |
| total FAST5 files | 18047 | 5058 | n.a. |
| total reads(2D,f,r) | 41305 | 15174 | 12309 |
| total base pairs | 226928730 | 92930784 | 83879212 |
| mean | 5493.98 | 6124.34 | 6814 |
| median | 5245 | 5874 | 6719 |
| min | 5 | 217 | 217 |
| max | 303643 | 36689 | 36689 |
| N25 | 9665 | 9893 | 10160 |
| N50 | 7457 | 7699 | 8082 |
| N75 | 5316 | 5540 | 6217 |
| Useable type 2D | n.a. | n.a. | n.a. |

Statistics output of Run 3 **FF**
| Input DATA | FAST5 FF | FAST5 FF CF >5kb < 29kb |
| --- | --- | --- |
| total FAST5 files | 12989 | 5152 |
| total reads(2D,f,r) | 26131 | 11317 |
| total base pairs | 133997946 | 92747729 |
| mean | 5127 | 8195 |
| median | 4834 | 7688 |
| min | 5 | 5023 |
| max | 303643 | 28185 |
| N25 | 9501 | 10218 |
| N50 | 7303 | 8332 |
| N75 | 5168 | 6869 |
| Useable type 2D | 5645 | 2940 |

The library was eluted and QC resulted in 38.6 ng/µl PSM concentration. Total mass 1003 ng. 6 µl were used as input for the SM. Flowcell QC: 380 active pores

**Figure 30.**
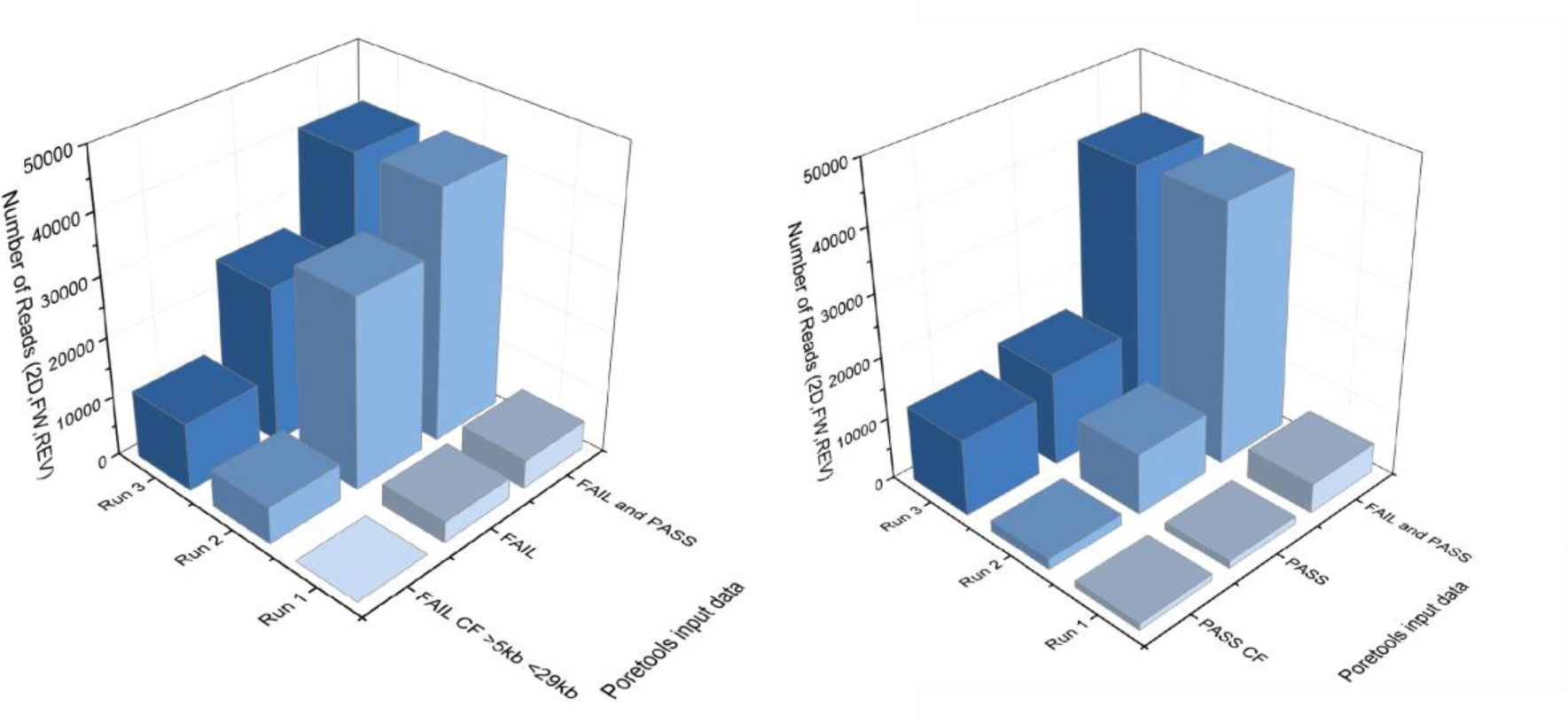
Corresponding 3D diagram to tables 8-10. Left: Data visualization corresponding to the fail filter (FF) of all three runs. Right: Data visualization corresponding to the pass filter (PF) of all three runs.

All 3 runs combined resulted in overall 44 942 FAST5 2D basecalled reads (FF and PF combined), an equivalent to 53.6 Gigabyte data which were basecalled successfully. A Total of 8 784 were 2D “high quality Metrichor^43^” PF basecalled FAST5 reads with an average readlength of 5308 bases. Resulting in theoretically 46m basepairs PF high quality data. The longest 2D basecalled consensus sequence was 36 kb long.

The following two pages contain the cumulative read count over sequencing time shown in figure 31 and the readlength distribution after removal of the control DNA reads shown in figure 32. This can be compared with control DNA included read length distribution shown in appendix figure 41.

**Figure 31.**
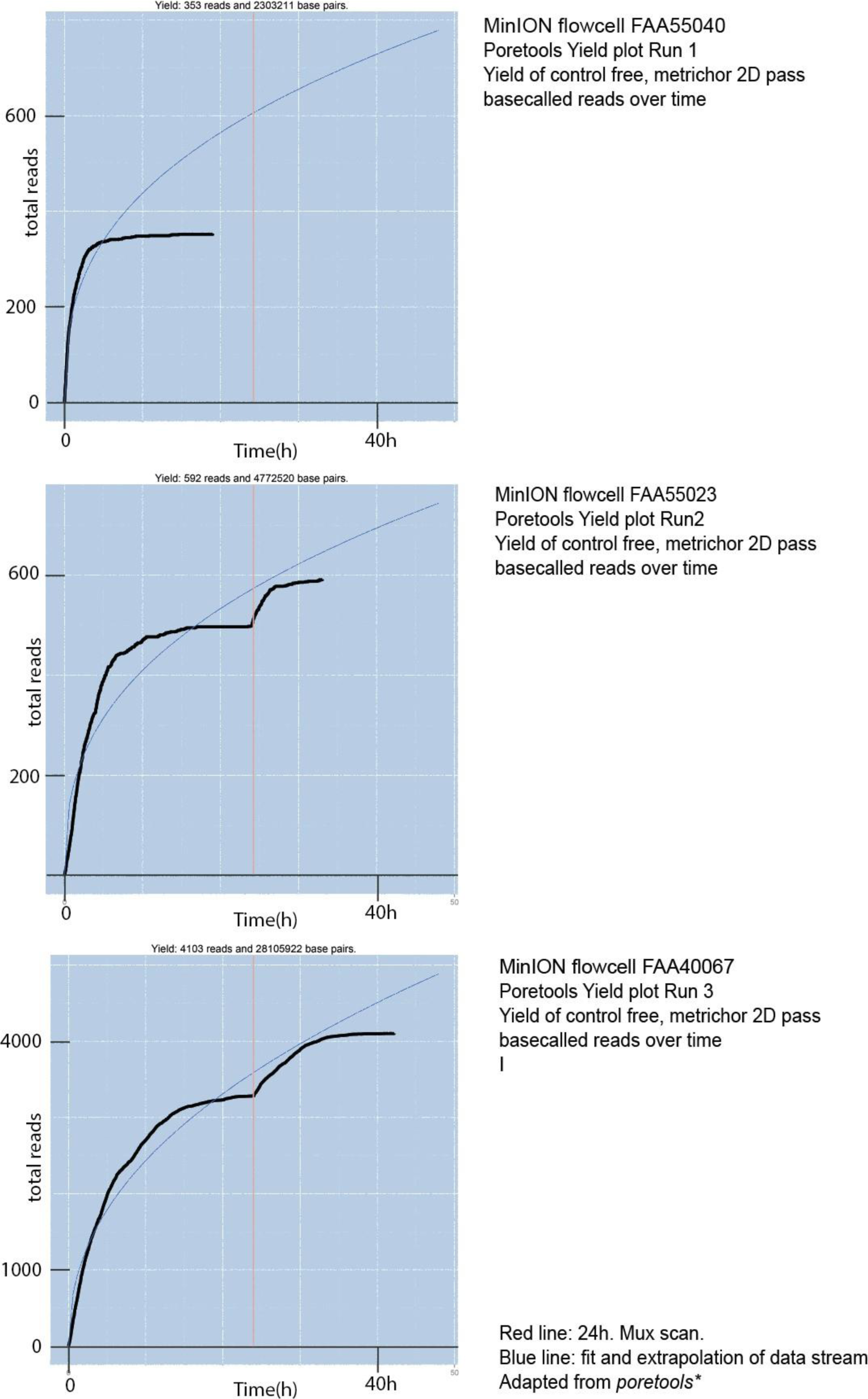
Yield Plot of Metrichor 2D pass basecalled and control free high quality reads. Adapted from R poretools^60^ The blue line represents a theoretic extrapolation. The Red line represents a “rescanning” of the nanopore sensors by the MinKNOW software (Mux scan).

**Figure 32.**
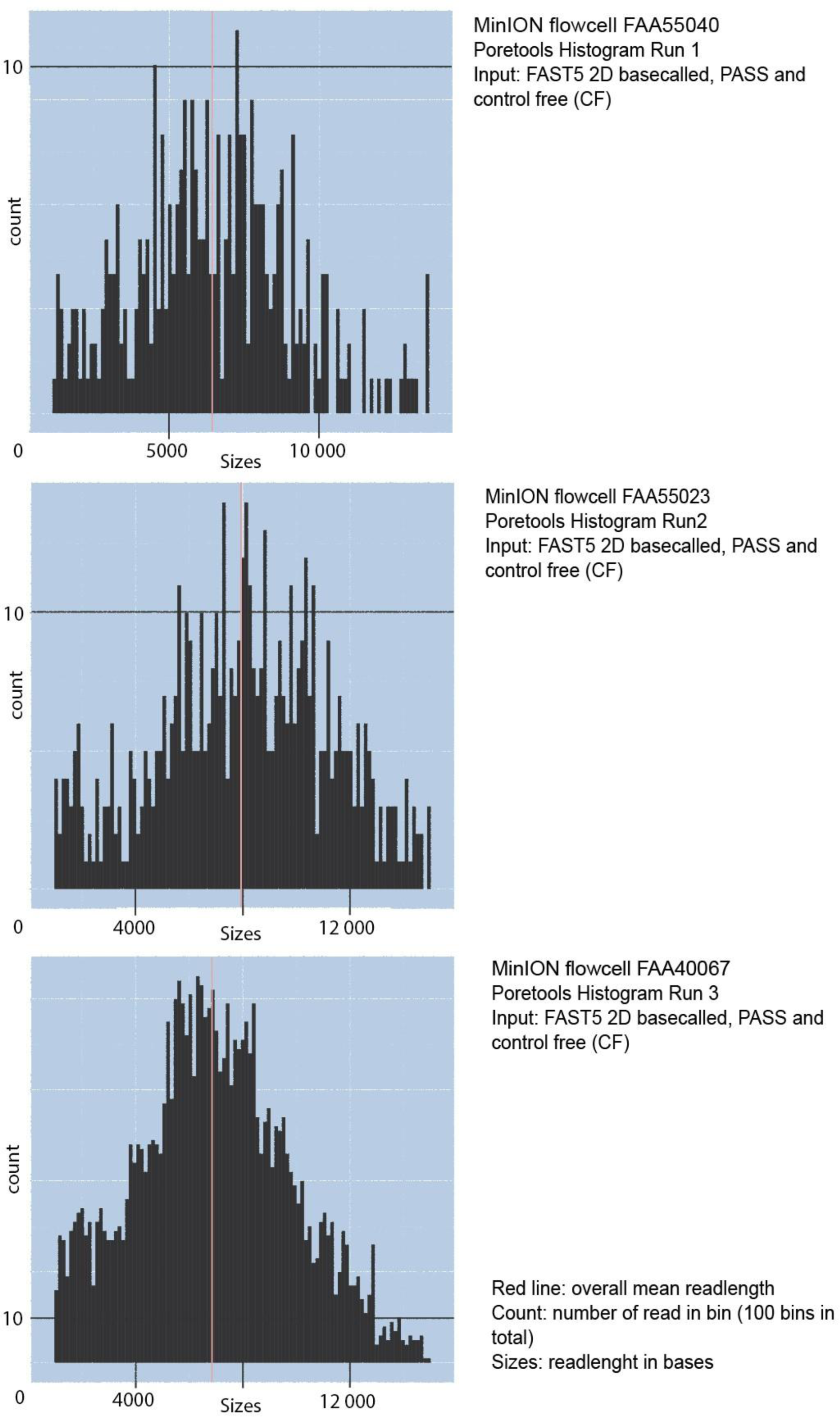
Histogram of Metrichor 2D pass basecalled and control free high quality reads. Adapted from R poretools. The red line shows the mean read length.

### 5.5 Genome assembly

After running the Metrichor basecalling service the raw reads were extracted and subsequently assembled with bioinformatics tools made available and described by Jared Simpson, Nikolas Loman and others. Compromising Poretools^60^, Mirabait^61^, Nanocorrect^2^ and Nanopolish^2^ in combination with the Celera^19^ assembler. The different assembly strategies and workflows are shown in figure 34. The results corresponding to the workflows are listed in figure 35/36. The different approaches resulted in different assemblies which are presented in the following tables. The assemblies vary depending on the coverage, quality and length of the input reads.

**Figure 34.**
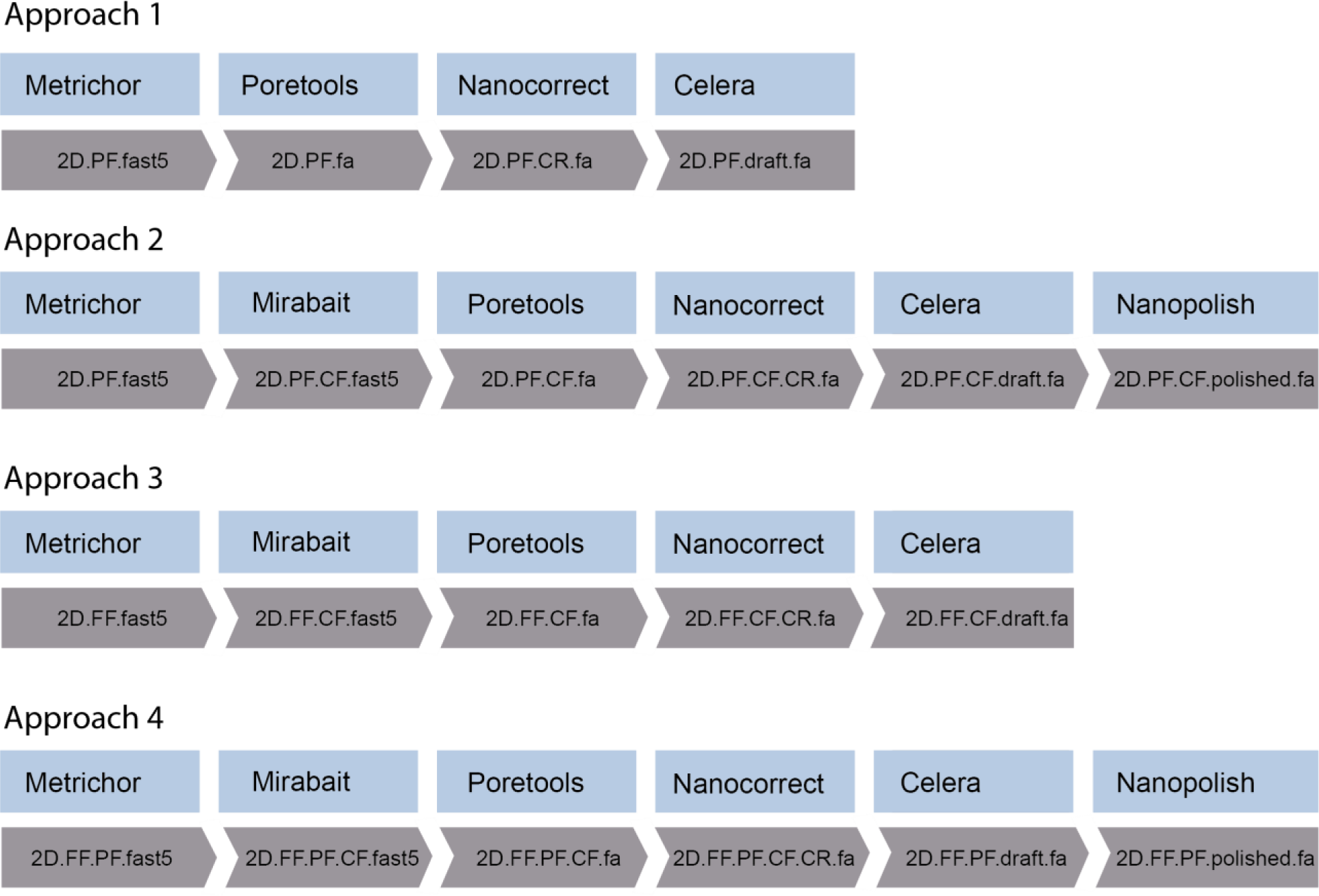
Assembly strategies corresponding to the results in figure 35 / 36.

**Figure 34.**
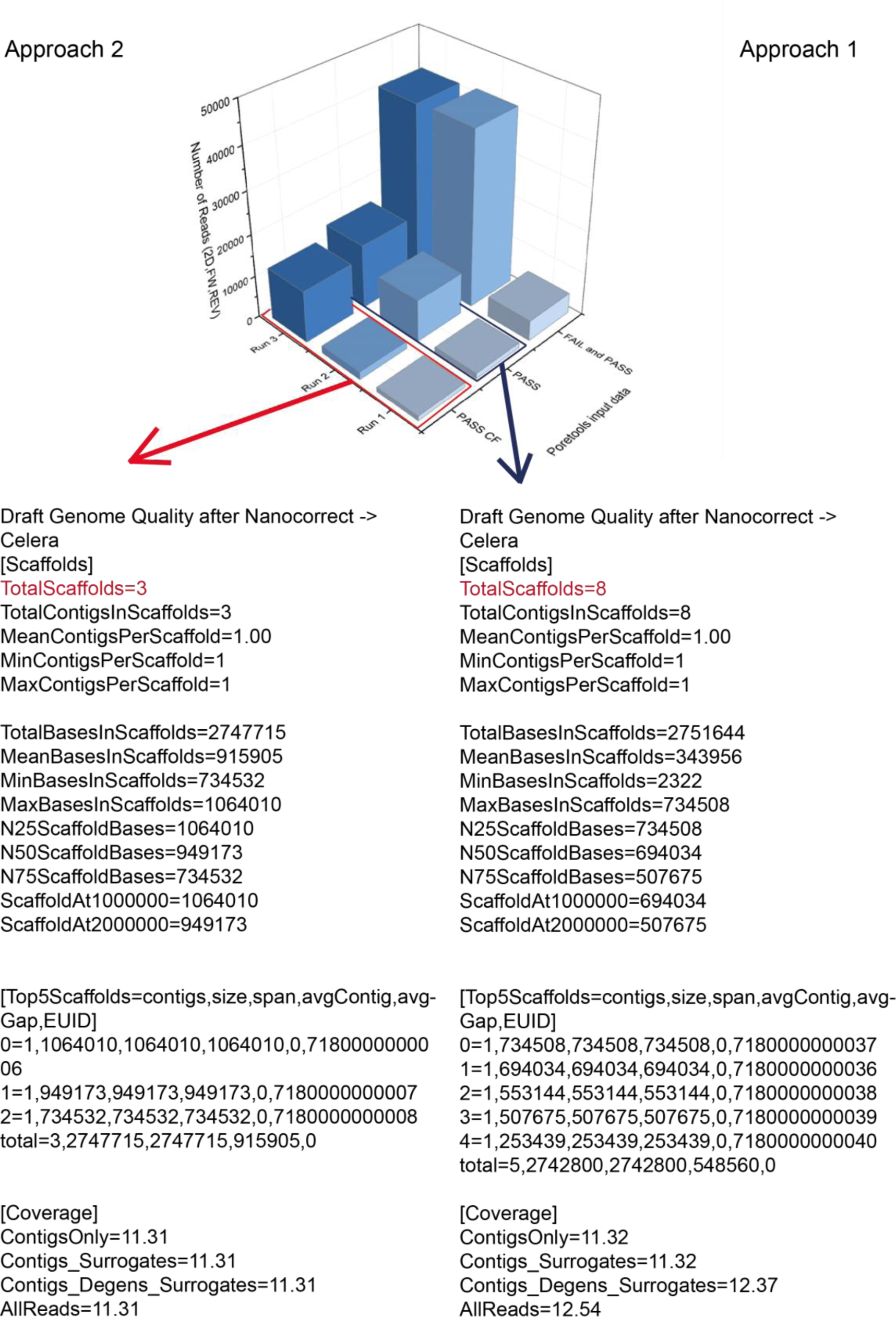
Quality report of the Celera assemblies for approach “1”(right) and “2”(left). The red/blue arrow refers to the data pool used for the assemblies.

**Figure 35.**
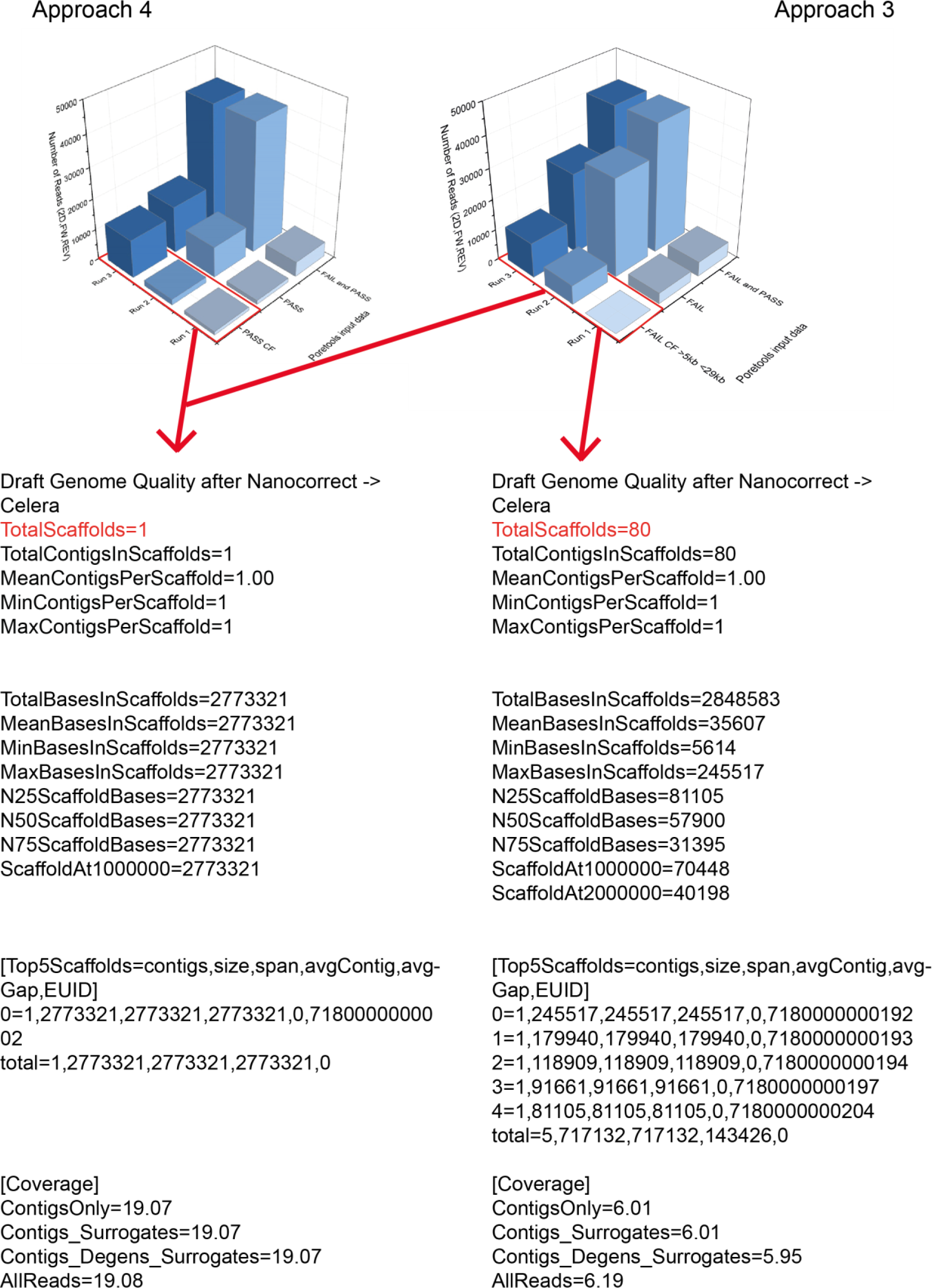
Quality report of the Celera assemblies for approach “3”(right) and “4”(left). The red arrow refers to the data pool used for the assemblies.

The following assembly results for “Approach 1, 2, 3 and 4” are shown.

In figure 34 the result for approach “1” and “2” are presented. In approach “2” an initial data prepossessing step was added to the Loman et al. workflow. To reduce noise during the assembly process, the internal control DNAcs reads were removed from the raw data.

The final draft genome correction with nanopolish did alter 3.1 % of all bases for approach “2”. The two corrected draft genomes (approach “2” and “4”) were compared by whole genome alignment, using LASTZ^65^ implemented in Geneoius© 9.0.4, to show consistent performance of the data. The whole genome alignments showed an average identity of 99.0 % over 2.8 megabases (mb). The step length of 500 000 and seed pattern 14 of 22.

### 5.6 Genome annotation

After the successful assembly, the genome was annotated with Prokka. A circular genome map was created by DNAPlotter^66^ shown in figure 36. The circularity has not been proved. A linear map showing important genes is included in the appendix. The map visualizes the final assembly with corresponding Prokka annotations. The annotation of the novel genome resulted in 7111 possible open reading frames. 2572 predicted proteins were assigned to these open reading frames.

**Figure 36.**
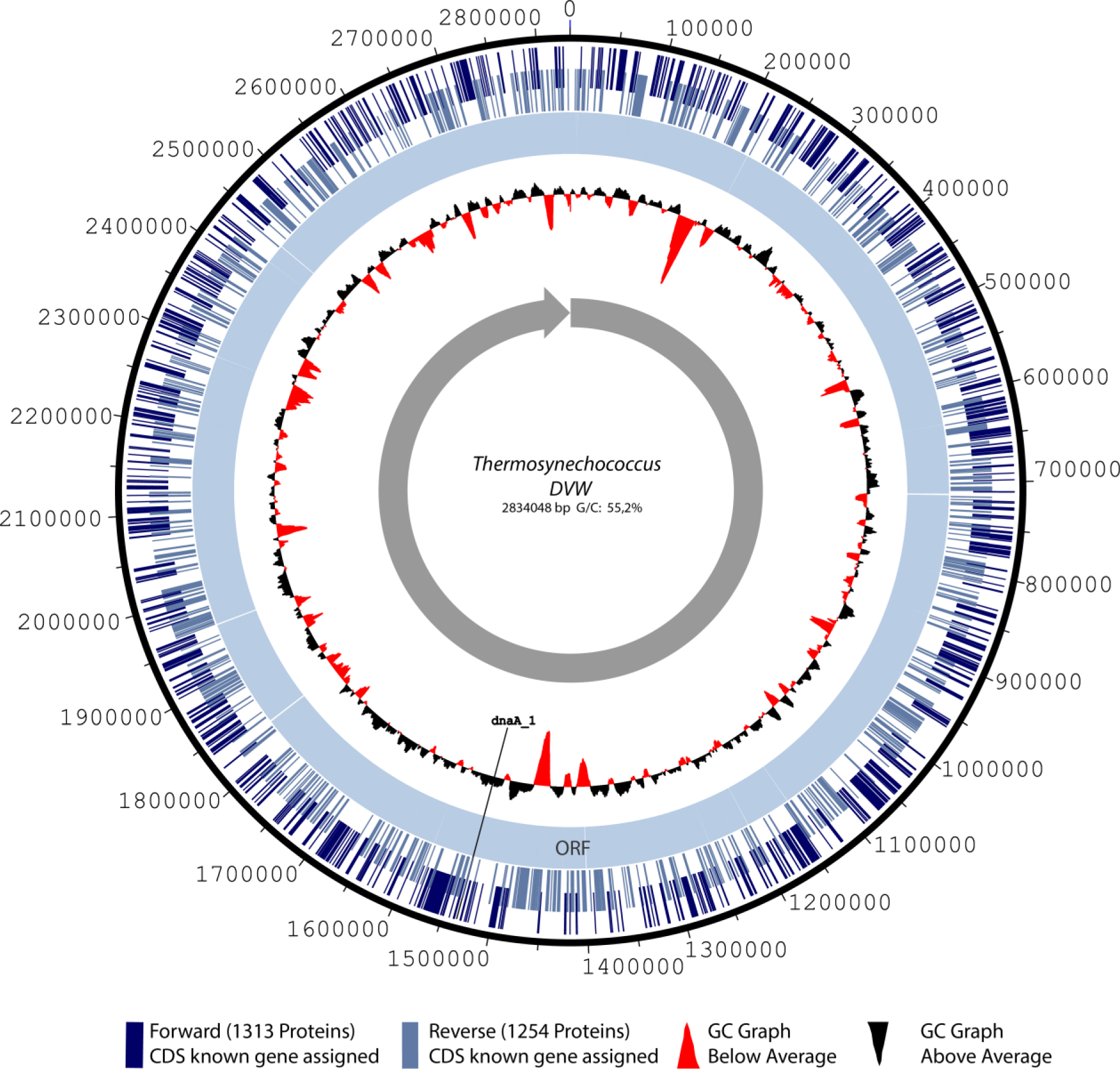
Circular genome map showing the position and orientation of predicted known genes, repetitive sequences and open reading frames (ORF/solid light blue circle). The outer dark blue circle represents the forward orientated genes. The outer light blue circle represents the reverse orientated genes. The inner black/red circle displays the GC content. *dnaA_1* marks the possible origin of replication. Assemblers like Celera do not suggest circularity, this has not been proven by further sequence mapping or primer walking.

The origin of replication is mostly in proximity of the *dnaA* gene shown in figure 37.^67^ The name of the substrain is provisional, a final proposal will be made during peer reviewed publication.

**Figure 37.**
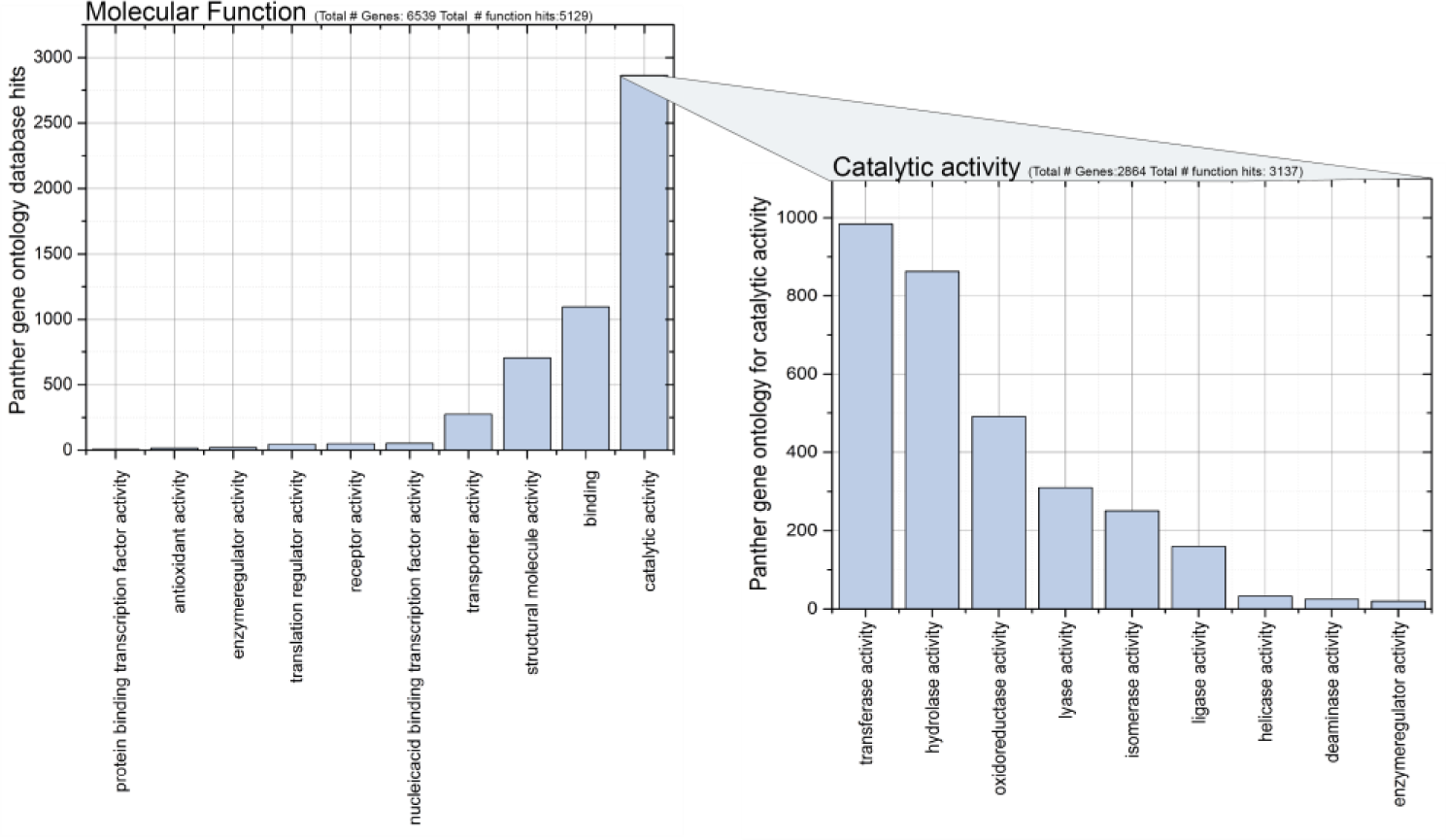
Gene ontology results from PANTHER^68^. Left: Molecular function classification by a subset of 5129 hits. Right: Subset of 2864 genes classified by their catalytic functions.

As part of the annotation process a gene ontology characterisation of the novel genome was conducted. The PANTHER (Protein ANalysis THrough Evolutionary Relationships)^68^ gene classification database was used to categorize the set of predicted genes. The PANTHER results for molecular function classification is presented in figure 37.

Further analysis showed that no coding sequence for plastocyanin (*PetE*) was identified by Prokka. It was observed that in the thermosynechococcus genus the gene is missing although it is regarded as part of “the cyanobacterial core genome”^69^.

Assembly approach 2 resulted in 2388 by Prokka predicted genes. The comparison of the annotation results of both assemblies “2” and “4” showed that 1753 genes (∼70%) were identical to the 2527 predicted genes of approach 4.

## Discussion

Starting with *Thermosynechococcus DVW*, more light will be shed on this new cyanobacteria member in future research projects.

A lot of effort during this project has been put into the MinION™ sequencing and *de novo* genome assembly. The MinION™ performance, the assembly quality and annotation results will be discussed later on. Some conclusive words on the overall costs, and future applications in the biotechnology industry, are presented at the end of this study.

### The novel *Thermosynechococcus “DVW”*

The 16s rRNA sequence BLAST^56^ (NCBI) results proved that the isolated wild type cyanobacterial strain is a member of the *Synechococcus* genus. Considerable genetic variation compared to the current database entries has been detected.

The 16s rRNA study by PCR proved that the enriched culture composition was mainly formed by *Synechococcus* subspecies. As the 16s rRNA PCRs were conducted before clone enrichment, the obtained sequence may be a result of superimpositions or a competitive PCR amplification. The final 16s rRNA study conducted with the genomic sequence extract, resulted in higher sequence similarity to known thermophilic strains. The first hit *Synechococcus lividus* in the NCBI BLAST results table 7 is a thermophile *Synechococcus* strain isolated by J.J. Copeland 1936. The “*Thermosynechococcus DVW*” strain was isolated from a hot spring and it is postulated that this strain belongs to the thermosynechococcus group introduced by Katoh et al. This assignment is supported by the finding that plastocyanin does not occur in *Thermosynechococcus*, and was also not predicted to the *Thermosynechococcus DVW* genome.

Furthermore, *Thermosynechococcus elongatus/NK55* does occur on the BLAST^56^ hit table 7. The novel genome is significantly (∼200kb) bigger than the two sequenced thermosynechococcus candidates. An additional comparison (not presented), of the novel genome by whole genome alignment, showed that there is significant evolutionary distance between *T. elongatus/Nk55* and *T. DVW*. This underlines the novel character of the strain which will be examined during further research.

### DNA sequencing with the MinION™

DNA sequencing with the MinION™ has some theoretical advantages which makes it an attractive DNA sequencing method, potentially reaching the $1000 genome aim^25^: The technology could be very cheap, being independent from special reagents, expensive laboratory space and power supply. The technology can be adapted to various adjacent fields like native RNA and protein sequencing.^25^ Direct detection of modified/methylated DNA bases seems to be feasible in the near future.

The genomic DNA fragments may be very long as MinION™ reads are currently spanning up to 200,000 bp.^20^ This is a major computational advantage to genome assembly^17^ (especially concerning repetitive regions). Moreover, the data can be accessed in real-time, making it an online analytics device.^1^

There are limitations of the technology: the data is a result of “stochastic sensing”^70^, making it to a certain degree, error prone. A high sample purity is necessary and homopolymer regions are less sensitively recorded than by other NGS methods. The currently used ASIC sensors are expensive and limited amount of data interpretation software is available. The “cloud” based basecalling service has opened discussion on data security risks.

Despite its commercial availability the device and its technology is still in a developmental stage. During this project it has been shown that the MinION™ can be run by a single person requiring little experience with next-generation sequencing devices. In the following, the MinION™ sequencing performance observed during this work is discussed.

### Sequencing performance

The overall MinION™ performance did allow to *de novo* assemble a prokaryotic microorganism genome, making it suitable for routine work with *E. coli* or other important microorganisms used in the biotechnology industry.

The quality of the flowcell units delivered by ONT was consistent. Interestingly, the flowcell with the lowest active pores produced the highest data yield. In appendix figure 42 a heatmap visualizes single high performing pores. Next to “bad” nanopores, a lot of possible error and uncertainty can be brought into the system by library preparation alterations. In the MAP community (and publications^20^) it has often been reported, that steps were forgotten or avoidable “errors” were made during the library preparation. Therefore, once again, the checklist type protocol should be used to reduce library preparation protocol variations.

Run “3” is regarded as the best run in terms of overall 2D basecalling results. 28% of the reads from Run “3” were Metrichor 2D pass filter files (PF). The size distribution for the 8 kb protocol shown in figure 41 was a bit smaller than expected, with a maximum at 6 kb (PF). Run “1” produced a low overall data yield, although 692 active pores were recorded during the so-called “Mux scan”. Run “2” resulted in low 2D output, although it produced the highest yield. The run sequenced mostly control DNA (DCS 3.5 kb dsDNA) which is visualized in figure 41 in the appendix. The 20 kb gDNA fragments used as input for run “2” are regarded as long. The library required that a mean of 40 000 bases progresses intact through the pores. The results show that the library was probably fragmented during library preparation, although greatest care was taken to minimise fragmentation risk. Previously, a competition for ultra-long reads had been initiated by ONT, but as the performance decreased with ultra-long reads^20^, it is now recommended (for high data yield) to stay in the 8-10 kb fragmentation target. As the ONT protocols and the “chemistry” is under constant development, protocols will vary in future from the presented.^1^ There is ongoing research done by ONT, to increase current performance by a multitude.^1,25^. A special research focus is on automated library preparation, which could minimize the observed performance bias.^1^

By updating the “chemistry” concepts and sensors, a bright sequencing future can be imagined.^71^ Using two protein or DNA origami pores (one for each strand) or an engineered hybrid helicase-nanopore complex, could enable library preparation free native DNA sequencing. - The limit is still the nanopore.

**Figure 38.**
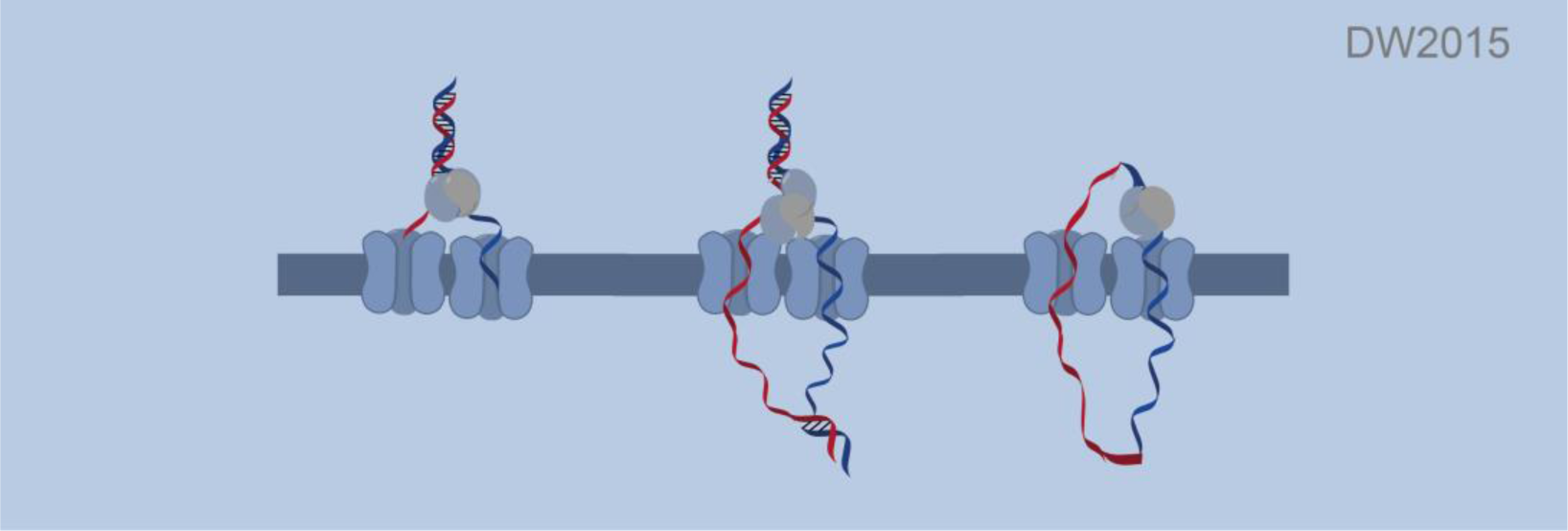
Imaginary concept of nanopore sequencing. Two single strands of the same DNA molecule propagating in a timed manner through two neighbouring pores.

### Assembly performance

Genome assembly was recently defined by Keith Bradmann (UC Davis) in the following way: “the art of trying to make one big thing from millions of very small things.” *De novo* assembly is still one of the most computational complex problems in biology.^17^ David Jaffe (Broad institute) pointed out that new technologies require new assembly methods, and therefore “every time the data changes, it is a new problem”.^17^ The long MinION™ reads are reducing the assembly complexity, but require significant read correction computational resources.

Unfortunately, the resulting assembly depends very much on the assemblers algortihm.^17^ As already pointed out in the introduction, there are many different assemblers to choose from. The comparison of genome assemblies is not made easier by the large diversity of algorithms used. There have been various studies trying to evaluate the assembler’s performance on a broad scale. The so-called Assemblathon and the Assembly Gold-Standard Evaluations (GAGE) should be pointed out here for further reading. It can be assumed that ONT will provide bioinformatics solutions which would increase comparability. A first attempt and realisation of this idea is WIMP, a Metrichor metagenomics workflow for a 16s rRNA based sample diversity estimation.^1^

The MinION™ gDNA reads were assembled to one continuous scaffold representing one bacterial chromosome. No additional plasmids were discovered, which is consistent to the observations from other strains of this group.^52^

The final circularisation of the genome was not shown and is still open for discussion. The comparison of the different assemblies presented in chapter 5.4 showed that the control DNA (DCS) has a negative impact on the overall assembly pipeline. These presented results had led ONT to include a filter to Metrichor, which will be available in future versions. Including the internal control during library preparation is not mandatory, but for the scenario in “Run 2” it was very useful for evaluation.

The assembly with FF and PF data combined resulted in the overall best assembly, in respect to scaffold number.

The scaffold number varied depending on input data amount and quality. The high quality data assembly “Approach 2” did not result in a continuous scaffold. This might be due to medium-low coverage, resulting in gap regions. The worst assembly resulted from approach “3” using low quality Metrichor 2D fail reads only. The finished assemblies of approach “2” and approach “4” were aligned by Geneious. The resulting overall identity of 99.0% does show that the MinION™ assembly process is still error prone. It seems that the mismatch regions are associated with homopolymer regions which is a known systematic error of MinION™ reads.^42^ During further work the different assemblies should always be compared. Further functional comparison of both assemblies will shed light on the overall correctness.

### Annotation of a *de novo* genome assembly

The annotation of *Thermosynechococcus DVW* resulted in 2572 predicted known genes. This number of predicted genes does correspond to what has been reported for *Thermosynechococcus elongatus BP-1*. The list of genes will be made available on Genbank in early 2016. In appendix figure 44, important genes corresponding to some “cyanobacterial core genome functions” are shown.^69^ Unfortunately, the assemblies are not identical and possible MinION™ related insertion or deletions (“indels”) are changing the reading frame, thus having great impact on annotation. The Prokka annotation results of the assembly approaches 2 and 4 were 70% identical. This shows that there are genes predicted to both assemblies, but many have to be checked by manual annotation. Reviewing these annotations by proteomics and other methods will be done during future research projects. The *PetE* gene was not predicted. This gene codes for plastocyanin, a protein which is transporting electrons to photosystem I.^72^ As this function is also not present in *T. elongatus BP-1* this might reveal a possible convergent evolutionary step. The gene ontology analysis by PANTHER^68^ shown in figure 37, visualizes that most genes fulfil catalytic functions. Within the catalytic functions, transferase activity is the most dominant group.

### Possible applications and cost of sequencing

Nothing was said up to now regarding the costs of nanopore sequencing. Such an essential question and driving factor of NGS devices, cannot be overlooked. The costs discussed in the following, are estimates and rely on personal experience. The presented calculations are only intended to provide an impression.

The initial cost of the device was $ 1250 paid to Oxford Nanopore Technologies™ as a MAP deposit fee. For every run the disposable consumables costs summed up to € 33. For DNA extraction and library preparation an additional € 52 were necessary. The final cost for a single library preparation sums to € 85. To this the cost, the capital for laboratory equipment needed should be added, including devices for sample preparation, storing and quality control.

“Do-it-at-home” DNA sequencing with the MinION™ is not yet possible, but the capital investments needed to conduct a sequencing run is extremely low compared to other sequencing platforms. We might be in the golden age of DNA sequencing^73^, but still it requires a lot of money to run experiments for routine applications.

As four flowcells were provided, a hypothetical run cost of € ∼400 is assumed in the following. By conservative calculation of “fast5 files” times “mean bp length”, an average (run 2&3) usable 96,742,981 bp translates into 241 kb / €. This results in an expensive ∼€ 13,000 for one haploid human genome (1x). Clearly, this is not the targeted price / performance range of ONT. Assuming that the MinION™ will reach the “$1000 Genome” aim, this would infer a tenfold price reduction. Achieving this goal would place the overall experiments cost in a range under € 100. This would possibly open doors to genome/transcriptome based screening applications as a routine method for bioprocess development. Moreover, the overall assembly process can be achieved by reasonable computational equipment for prokaryotes, requiring strong workstation PCs but not expensive computational clusters.

The importance and relevance of “personnel” sequencers to industrial biotechnology cannot be overemphasized. Integrity control after genomic integration experiments and gene expression levels comparisons are a few of many other possible applications.

### Outlook

*Thermosynechococcus DVW* belongs to a group of microorganisms responsible for global scale CO2 fixation. The cyanobacteria phylum has been gaining popularity for possible energy harvesting and pharming concepts. Further research focussing on this strain will follow in the near future. Nanopore sequencing by the MinION™ device is sufficiently accurate to sequence prokaryotes. As the technology improves, larger eukaryotes will probably be sequenced in the near future using one flowcell only. Oxford Nanopore Technologies™ announced that new product iterations will be available to MAP participants, thereby generating high hopes of reaching new horizons with MinION™ sequencing. Nanopore DNA sequencing is, despite current performance drawbacks, viable in every standard laboratory setting, making genomics research more independent than ever. As already pointed out, the technology has room for various further improvements. Only the imagination can set the limits to the application spectrum.

## Acknowledgments

Many thanks to the Oxford Nanopore Technologies™ team who were extremely motivated, kind and helpful. We would like to give thanks to all partner departments for providing free access to analytical devices, including Dr. Wenning (Qubit at ZIEL II MySeq Lab), Dr. Haslbeck (Nanodrop photometer Department Biotechnology), Prof. Berensmeier (providing separation magnets), and Dr. Machuy (sponsoring required Ampure beads (Beckman Coulter).

